# Object representations drive emotion schemas across a large and diverse set of daily-life scenes

**DOI:** 10.1101/2025.02.19.638854

**Authors:** Chuanji Gao, Susan Ajith, Marius V. Peelen

## Abstract

The rapid emotional evaluation of objects and events is essential in daily life. While visual scenes reliably evoke emotions, it remains unclear whether emotion schemas evoked by daily-life scenes depend on object processing systems or are extracted independently. To explore this, we collected emotion ratings for 4913 daily-life scenes from 300 participants, and predicted these ratings from representations in deep neural networks and fMRI activity patterns in visual cortex. AlexNet, an object-based model, outperformed EmoNet, an emotion-based model, in predicting emotion ratings for everyday scenes, while EmoNet excelled for explicitly evocative stimuli. Emotion information was processed hierarchically within the object recognition system, consistent with the visual cortex’s organization. Activity patterns in the lateral occipital complex (LOC), an object-selective region, reliably predicted emotion ratings and outperformed other visual regions. These findings suggest that emotion processing in everyday scenes follows visual object recognition, with additional mechanisms engaged when object content is uninformative.

## Introduction

The rapid emotional evaluation of objects and events facilitates the pursuit of valuable resources and the avoidance of potential harm. Accordingly, visual scenes readily and reliably evoke feelings of happiness, disgust, awe, horror, amusement, etc. (Cowen & Keltner, 2017; Cowen et al., 2021; Ekman, 1992; Kragel et al., 2019; Lench et al., 2011; Riegel et al., 2016). Previous research has shown that there is agreement between observers in the emotion category labels they use to judge their emotional experience when viewing emotionally evocative stimuli (e.g., scenes showing births, risky stunts, sexual acts; (Cowen & Keltner, 2017; Kragel et al., 2019). For example, one study found that 2185 emotionally evocative short videos consistently elicited 27 distinct emotional categories, which provided a more accurate representation of emotional experiences than ratings on 14 affective dimensions, such as valence and arousal (Cowen & Keltner, 2017). These findings prompt further investigation into the neural and computational mechanisms underlying the evocation of diverse emotions by visual scenes.

One possibility is that emotions are inferred from the objects, object states, and/or object relations that are present in a scene. For example, a scene showing a person holding a gun could evoke feelings of fear, sadness, or horror, while a scene showing prepared food on a plate could evoke feelings of craving. On this account, emotional experience would follow the recognition of the visual scene, with the visual processing stage of the process relying on known mechanisms of object and scene recognition in ventral temporal cortex (Epstein & Baker, 2019; Grill-Spector et al., 2001; Grill-Spector & Malach, 2004; Malach et al., 1995; Peelen et al., 2024).

Alternatively, emotions may be evoked by visual cues before, or in parallel with, object recognition. Certain visual features could be consistently associated with specific emotions and evoke emotions without requiring object recognition (Bar & Neta, 2006; Lakens et al., 2013). Evidence for this account comes from a study showing that a convolutional neural network trained to categorize emotions from visual images (EmoNet) accurately predicts emotion ratings of human observers (Kragel et al., 2019). Importantly, EmoNet outperformed emotion categorization using object category labels from AlexNet, a convolutional neural network trained to categorize objects in scenes (Krizhevsky et al., 2012, 2017). EmoNet was created by keeping all layers of AlexNet fixed, except for the last fully connected layer, which was retrained. This adjustment shifted the focus from classifying images into 1000 object categories to classifying images into 20 emotion categories. The connections between sensory features and emotions are referred to as emotion schemas (Kragel et al., 2019). Furthermore, the study showed that patterns of visual cortical activity could be used to decode human emotion ratings. Together, these findings suggest that emotion may serve as an organizing principle of the visual system, with emotion schemas being extracted from visual input without being mediated by object recognition.

EmoNet was developed using an emotional scene database (Kragel et al., 2019) that included emotionally evocative stimuli gathered by querying search engines and content aggregation websites with contextual phrases targeting various emotion categories (Cowen & Keltner, 2017). While these stimuli encompassed a broad array of psychologically significant situations, they were explicitly selected emotional scenes that may not be representative of our daily-life experience. Furthermore, most of the scenes involved human actions, such that there was relatively little variability in terms of object content across the image set. This raises the possibility that the inferior performance of AlexNet (relative to EmoNet) reflected the lack of variability in the object categories present in the scenes. While human actions are clearly important for many expressive emotions, we also experience emotions for a wide variety of daily-life scenes that do not include humans. For example, images of food (associated with craving), spiders (associated with fear), or flowers (associated with aesthetic appreciation) are also consistently tied to specific emotions. This raises the question of whether an emotion-specific recognition mechanism, such as modelled by EmoNet, is also used for evaluating emotions from a broader and more representative set of scenes.

To address this question, the present study took a large and diverse set of images for which fMRI data are available (Chang et al., 2019). We presented this extensive collection of approximately 5000 images to 300 volunteers to gather emotion ratings for each image. Specifically, the dataset comprised 1000 images depicting indoor and outdoor scenes across 250 categories with a general focus rather than a focus on specific objects, actions, or people; it included 2000 complex images featuring multiple objects, often set within realistic contexts and depicting interactions with other animate or inanimate entities; and it included 1916 images predominantly showing individual objects, covering 958 distinct object categories. Analyzing this dataset allows for a comprehensive evaluation of the hypothesis that emotions are evoked through an emotion-specific recognition system rather than based on object representations. The fMRI dataset included data from participants who observed approximately 5000 unique images, which is more effective for uncovering universal principles of human brain function and offers several unique benefits over studies that involve a more restricted set of stimuli (Naselaris et al., 2021).

Our results demonstrate that while an emotion-driven model (EmoNet) excels in predicting emotion responses to explicitly evocative stimuli, an object-based model (AlexNet) better captures emotion responses to a broader set of common, everyday scenes. Additionally, we found that emotional information is represented hierarchically within the object recognition system, with the lateral occipital complex (LOC) playing a particularly prominent role. These findings indicate that emotions evoked by daily-life scenes are mediated by object recognition. Only when object content is not informative, additional visual processing mechanisms are needed to map visual input to emotional experience.

## Results

### Object representations in deep neural networks predict emotion ratings of daily-life scenes

Here, we tested whether emotion recognition relies on emotion-specific visual processing or can similarly be explained by established object and scene processing systems. To investigate this, we used the AlexNet deep convolutional neural network (DCNN) model as a representation of object processing (Krizhevsky et al., 2012, 2017). AlexNet was chosen due to its established use in vision research and to allow comparison with EmoNet (Kragel et al., 2019). EmoNet, a convolutional neural network derived from AlexNet, shifts its focus from object classification to categorizing images into 20 distinct emotion categories. We hypothesized that if emotion acts as a general organizing principle within the visual system, EmoNet should outperform AlexNet in predicting emotion ratings. Alternatively, if emotion recognition relies on existing object processing systems, AlexNet should be at least comparably effective in predicting emotion ratings relative to EmoNet.

We recruited 300 volunteers who viewed 4913 images for which fMRI data are available (BOLD5000; Chang et al., 2019). Three out of the 4916 unique images were not included in the experiment due to a technical issue. The 4913 images were randomly distributed into 30 sets: 29 sets contained 165 images each, while one set included 128 images (**Fig. 1a**). Volunteers were assigned to evaluate each set separately, with the goal of having 10 participants per set. The sample size aligns with prior studies demonstrating that judgements from approximately 10 participants are sufficient to reliably approximate population-level means (Cowen et al., 2018; Cowen & Keltner, 2017). After viewing each image, participants were asked to respond to a prompt featuring 20 emotion categories: adoration, aesthetic appreciation, amusement, anxiety, awe, boredom, confusion, craving, disgust, empathic pain, entrancement, excitement, fear, horror, interest, joy, romance, sadness, sexual desire, and surprise. We deliberately adopted the 20 emotion categories employed by Kragel et al. (2019) to enable a fair comparison of the predictive performances of EmoNet and AlexNet, as EmoNet is trained to classify these 20 emotion categories. Participants were instructed to select at least one category that best described their emotional response to the image, but they could choose multiple categories if desired (**Fig. 1b**). For each image, we recorded the frequency with which each emotion category was selected by the 10 participants. The probability of each of the 20 emotion categories was then calculated by dividing the number of times an emotion category was chosen by the total number of participants, which was 10 (**Fig. 1c and Fig. S1**).

**Fig. 1.**
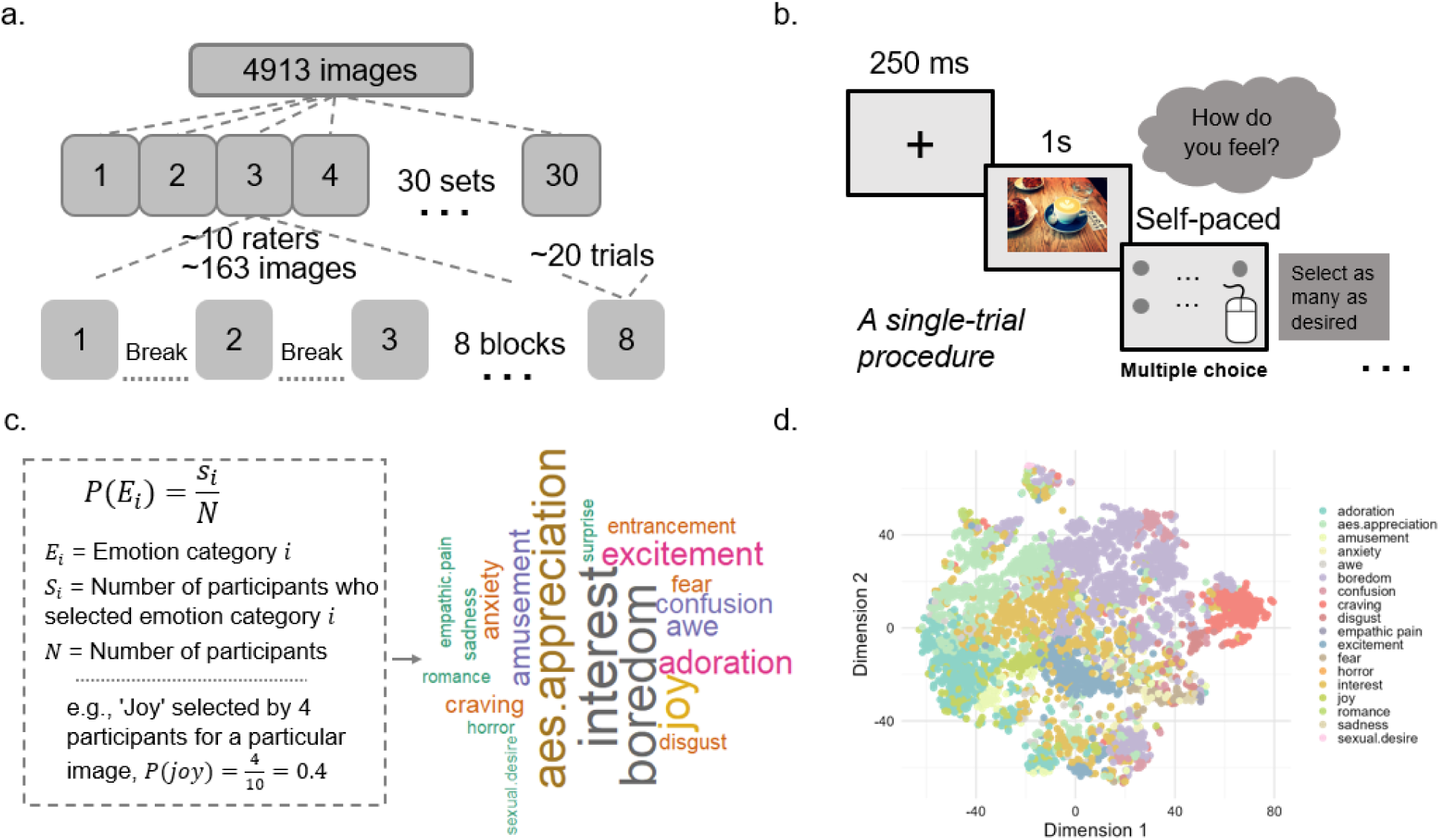
Behavioral study procedure and rating results. a) Overall experimental procedure. The 4913 images were randomly divided into 30 sets, with a target of 10 participants per set. Each participant rated around 163 images, which were divided into eight blocks. b) A single-trial procedure. Every trial began with a 250 ms fixation cross, succeeded by the presentation of an image for one second. This was followed by the emotion category prompt. If participants were uncertain about the meaning of any emotion while responding, they could view a description of each emotion by moving the mouse over the emotion category labels. c) Computation of emotion probabilities. The mean probability for each of the 20 emotion categories was calculated by dividing the number of times an emotion category was selected by the total number of participants, which is 10. d) Structure of images revealed by the t-Distributed stochastic neighbor embedding (t-SNE) analysis. Each dot was marked according to the emotion that had the highest probability.

We evaluated the reliability of emotion ratings for images by assessing the proportion of pictures exhibiting significant concordance in judgement rates across the 20 emotion categories following Cowen et al. (2017). We found that out of the 4913 images, 79% of the pictures have significant concordance (or rates of interrater agreement) for at least one category of emotion across raters [FDR < 0.05]. 55% of the raters chose the most agreed-upon emotion category for each picture [chance level = 13.5%, Monte Carlo simulation of all category judgments matching the same overall proportions of categories that were selected by the real participants]. These results are comparable or better to those documented in previous studies (e.g., Cowen et al., 2017) (**Supplementary text**).

We explored the structure of images using the technique of t-Distributed stochastic neighbor embedding (t-SNE). The t-SNE analysis revealed that images were distinctly clustered within the two lower-dimensional spaces (**Fig. 1d**). Each dot was color coded based on the emotion with the highest probability, showing that images of the same emotion category are more likely to cluster together. The t-SNE analysis was applied to the full 20-dimensional emotion probability vectors (derived from participant ratings) for each image, rather than relying solely on dominant emotion labels. This method preserves the complete distribution of emotion responses, including secondary and tertiary emotional associations. The resulting grouping of images by emotion category in a lower-dimensional space reveals inherent structural patterns in the data that extend beyond dominant labels. Furthermore, the clustering of images by their dominant emotion underscores the robustness and consistency of primary emotion responses as a key organizational feature of the dataset. Emotion dissimilarity between each pair of emotions and hierarchical clustering analyses showed meaningful clustering of emotion categories (**Supplementary text and Fig. S2**).

Having established that common everyday scenes and objects we frequently encounter are associated with various emotions, we proceeded to address our primary research question: comparing the performance of AlexNet and EmoNet in predicting emotion ratings. EmoNet differs from AlexNet in the retraining of the weights in the final fully connected layer (fc8), while the preceding seven layers are identical between the two models. Therefore, we focused our comparison on the predictive performance of the fc8 layer in both models in relation to emotion ratings. We used the partial least squares regression (PLSR) analysis approach (Abdi, 2010; Krishnan et al., 2011). This multivariate, data-driven approach identifies latent variables within a multidimensional input space (fc8 layer activations) and a multidimensional output space (emotional ratings) that are optimized to maximize the covariance between the two variable sets, enabling the prediction of emotion ratings from deep neural network layer activations (**Fig. 2a**). The choice of PLSR was motivated by three main reasons: first, its efficacy in handling datasets characterized by predictors that exhibit both high multicollinearity and high dimensionality; second, its capability to model complex multivariate patterns across both predictors and outcomes; and third, it is particularly suited for predicting behavior at the item-level, which is our primary objective.

**Fig. 2.**
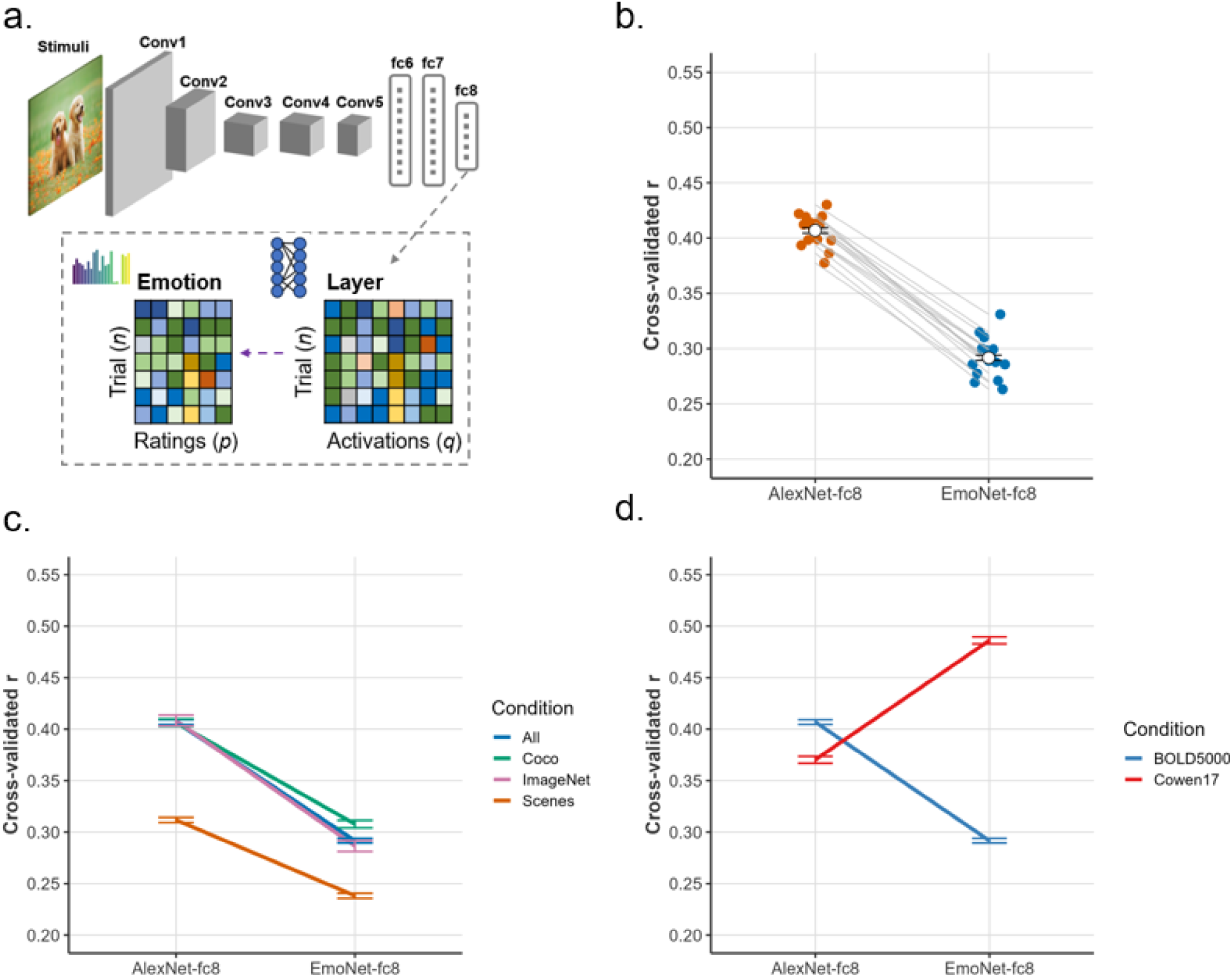
Object representations predict emotion ratings of daily-life scenes. a) Decoding emotions from deep convolutional neural network (DCNN) representations using partial least squares regression. b) fc8 layer in the AlexNet model outperformed fc8 layer in EmoNet model in predicting emotion ratings. Each line and dot represent the result of a cross-validated fold. c) AlexNet consistently outperformed EmoNet in predicting emotion ratings across three subsets of BOLD5000 images. d) EmoNet outperformed AlexNet in predicting emotion ratings for the Cowen17 dataset. In contrast, for the BOLD5000 dataset, AlexNet outperformed EmoNet in predicting emotion ratings. Error bars represent the standard error across cross-validated folds.

We found that both fc8 layer of AlexNet (average leave-one-session-out cross-validated *r* = 0.407, *p* < 0.001, permutation test) and fc8 layer of EmoNet (average leave-one-session-out cross-validated *r* = 0.292, *p* < 0.001, permutation test) significantly predicted emotion ratings. The confusion matrices demonstrated no bias in predictions (**Fig. S3**). Importantly, AlexNet outperformed EmoNet in predicting emotion ratings (Δ*r* = 0.115, *p* < 0.001, permutation test) (**Fig. 2b**). The top example images predicted from the fc8 layer of AlexNet for each emotion align with intuition. For instance, images of mountains, flowers, and houses are associated with aesthetic appreciation; pizza, dishes, and other foods are linked to craving; cockroaches, tiger beetles, and other insects are related to horror; bulletproof vests and military uniforms are associated with sadness; while faces, sunglasses, and women wearing lipstick are connected to sexual desire (**Fig. 3 and Fig. S4**). These examples show that common everyday scenes and objects are consistently tied to different emotions.

**Fig. 3.**
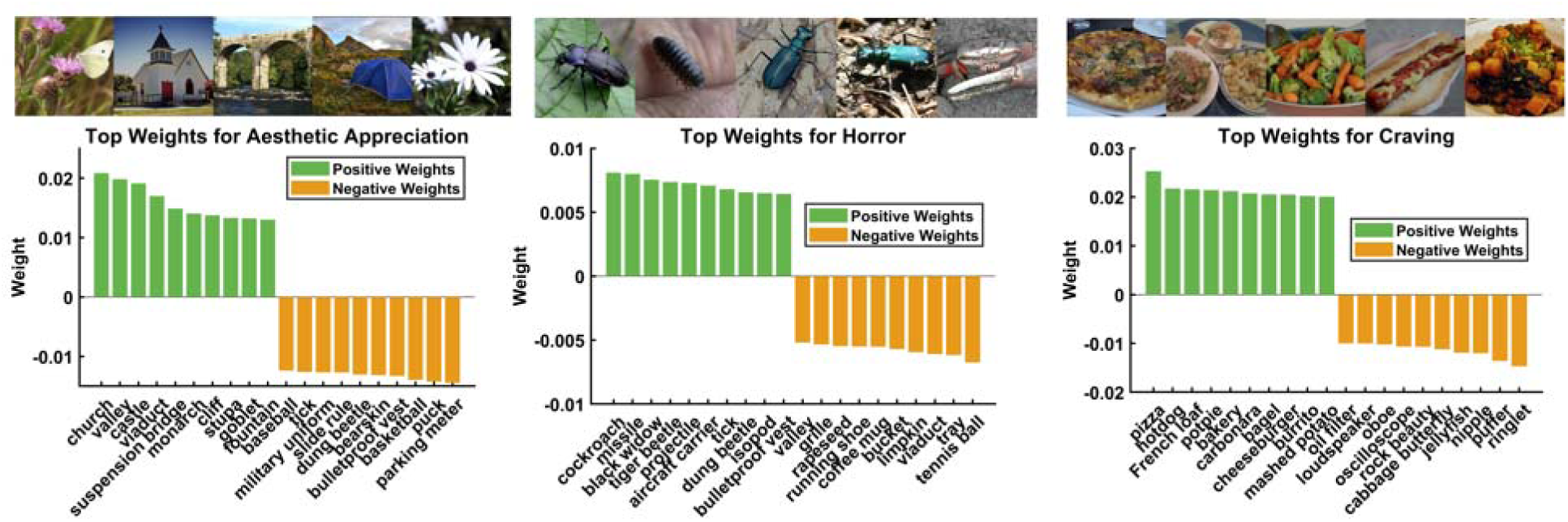
Top 5 example images predicted from the fc8 layer of AlexNet and the top weights for 3 example emotions. We present the top-10 positive weights and the top-10 negative weights. The 1000 (object) x 20 (emotion) weight matrix was derived by averaging the weight matrices from 15 cross-validation folds.

To evaluate the consistency of the comparison between AlexNet and EmoNet across diverse image datasets, we analyzed three subsets of BOLD5000 images: Scenes, COCO (Lin et al., 2014), and ImageNet (Russakovsky et al., 2015). The Scenes subset consists of 1000 images depicting both indoor (e.g., restaurants) and outdoor (e.g., mountains and rivers) environments, with a general focus on the broader scene rather than specific objects, actions, or people. The images encompassed 250 unique scene categories, primarily drawn from the SUN dataset (Xiao et al., 2010), with images selected using Google Search queries based on the category names. The COCO subset includes 2000 complex images from the COCO dataset, featuring multiple objects, often situated in realistic contexts and involved in interactions with other animate or inanimate entities (e.g., scenes of human social interactions). The ImageNet subset comprises 1916 images, predominantly depicting individual objects, selected from the ImageNet dataset. These three subsets are distinct from one another, and comparing the results across them allows us to assess the consistency of our findings across the broader set of images in BOLD5000. The results consistently show that AlexNet outperformed EmoNet in predicting emotion ratings (**Fig. 2c**).

In addition, given that a previous study (Kragel et al., 2019) demonstrated that EmoNet outperformed AlexNet in predicting emotions, we analyzed the images from Cowen and Keltner (2017) used in that study to assess whether we could replicate their findings and to examine differences in results between the BOLD5000 images and this earlier dataset. From each video in the Cowen17 dataset, we extracted three representative frames at 25%, 50%, and 75% of the video length, yielding 6555 images. Unlike the BOLD5000 images, which depict common everyday scenes and objects that we frequently encounter, the Cowen17 images consist of more emotionally evocative stimuli that were explicitly selected for their emotional content. We found that EmoNet outperformed AlexNet in predicting emotion ratings for these images, *t*(14) = 24.55, *p* < 0.001, paired t-test (**Fig. 2d**), which is consistent with Kragel et al. (2019). This finding contrasts with the results from BOLD5000 images, where AlexNet outperformed EmoNet in predicting emotion ratings. For explicitly emotionally evocative scenes, EmoNet (an emotion-based visual system model) outperformed AlexNet (an object-based visual system model), while AlexNet was superior for common daily-life scenes.

### Emotion information is processed in hierarchical stages of the object recognition system

Having established that object representations reliably predict emotion ratings of daily-life scenes, we next aimed to determine whether emotional information is processed in a hierarchical manner. If emotion recognition is mediated by known object processing systems, as we have demonstrated, we would expect emotional information to be processed hierarchically, consistent with the organization of the visual cortex for object recognition. AlexNet comprises five convolutional layers and three fully connected layers, mirroring the hierarchical structure of regions in the ventral visual stream (Eickenberg et al., 2017; Güçlü & Van Gerven, 2015; Khaligh-Razavi & Kriegeskorte, 2014; Wen et al., 2018). As activation progresses through the layers, the processed features grow increasingly complex, starting from low-level features in conv1 to complex object parts in conv5. Each convolutional neuron is connected to a limited subset of neurons in the next convolutional layer, transferring only a portion of the top-weighted activations to the next layer. Information from conv5 is then passed to the fully connected layers (fc6-8), where each neuron connects with all neurons in the next fully connected layer.

We employed PLSR to explore the relationship between layer activations and the emotion ratings. Specifically, we predicted emotion probabilities based on activations from various layers of AlexNet, including conv1, conv2, through to fc8. We found that emotion ratings could be predicted above chance from activity in all layers (*p*s < 0.001, permutation test, FDR corrected; **Fig. 4a; Supplementary text and Fig. S5**), and confusion matrices revealed no evidence of systematic bias in predictions (**Fig. S3**).

**Fig. 4.**
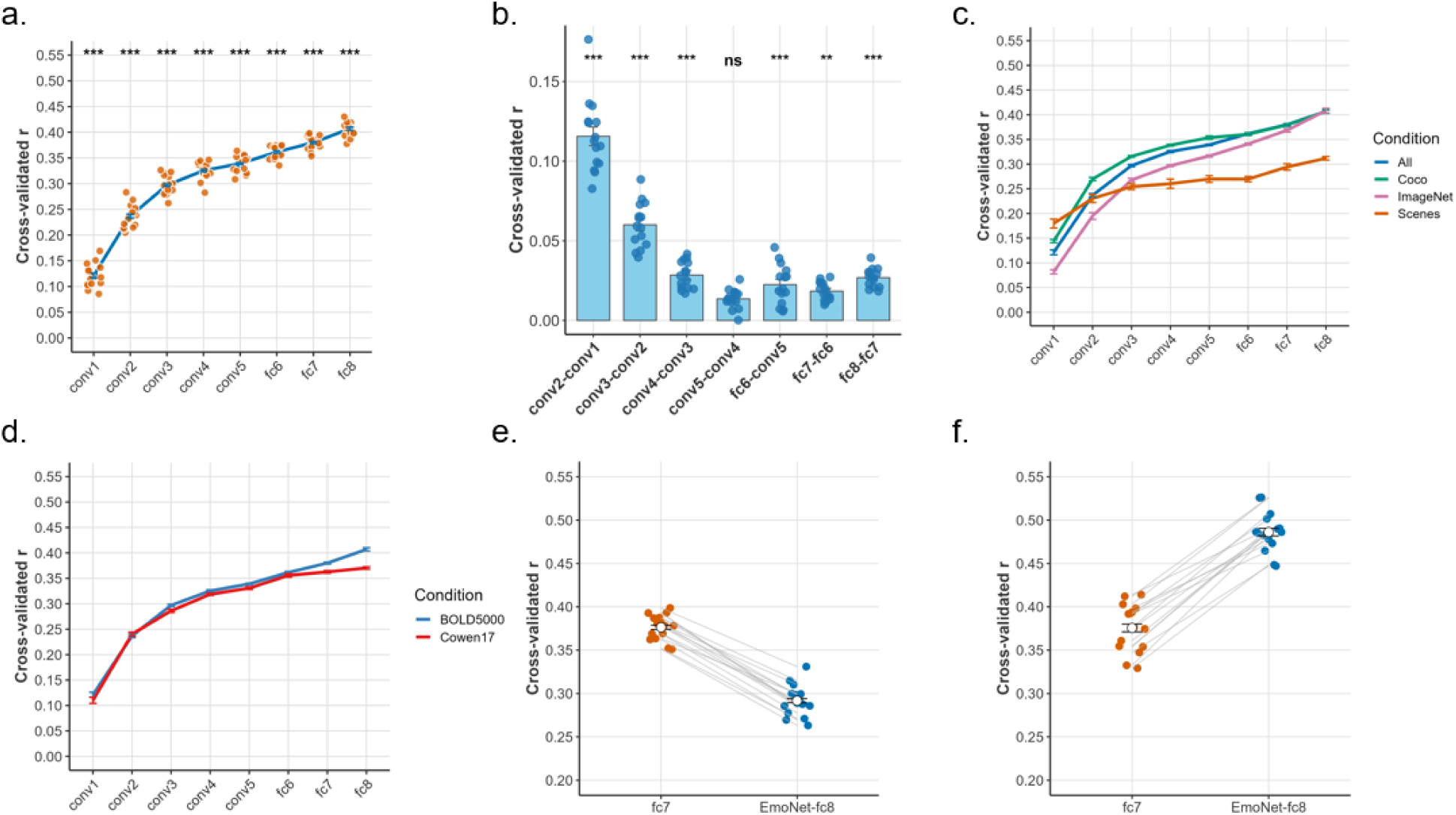
Emotion information is hierarchically represented in an object recognition system. a) Results of decoding emotions from AlexNet representations. These results show that emotion information is processed hierarchically in a visual object processing system. b) Differences between AlexNet layers in predicting emotion ratings. c) The hierarchical processing of emotion information was consistent across the three subsets of BOLD5000 images. d) The hierarchical processing of emotion information was consistent regardless of whether the BOLD5000 or Cowen17 dataset was used. e) fc7 layer outperformed fc8 layer of EmoNet in predicting emotion ratings for the BOLD5000 dataset. f) fc8 layer of EmoNet outperformed fc7 layer in predicting emotion ratings for the Cowen17 dataset. Error bars represent the standard error across cross-validated folds and each dot represents the result of a cross-validated fold.

To compare the difference across layers (conv1, conv2, conv3, conv4, conv5, fc6, fc7, and fc8), we performed repeated measures ANOVAs on the prediction-outcome correlation across different cross-validation folds, and Greenhouse-Geisser corrections were applied where necessary to account for violations of sphericity. We found a main effect of layer: *F*(2.93, 41.04) = 999.42, *p* < 0.001, partial = 0.99. These results demonstrated that different layers vary in their capability to predict emotion ratings (**Fig. 4a**). In addition, we examined whether emotional information is processed in hierarchical stages. Results showed significant improvements across successive layers, suggesting a monotonic increase in predictive power with layer depth (*p*s < 0.001, permutation test, FDR corrected; **Fig. 4a-b; Supplementary text**).

To evaluate the layered processing results across diverse image datasets, we analyzed three subsets of BOLD5000 images: Scenes, COCO, and ImageNet. We observed hierarchical processing of emotional information consistently across the three subsets of BOLD5000 images (**Fig. 4c**). In addition, we examined the layered processing results for the Cowen17 stimuli set. The results consistently show that deeper layers yield better predictions of emotional ratings compared to earlier layers in AlexNet representations, regardless of whether the BOLD5000 or Cowen17 dataset was used (**Fig. 4d**). These findings suggest that the hierarchical nature of emotional processing within the object recognition system remains consistent across different stimuli. However, whether emotion recognition primarily depends on the established object recognition system may vary depending on the specific stimuli.

We evaluated the predictive performance of layer 7 (fc7) and EmoNet layer 8 (fc8) activations for emotion ratings with the two datasets: BOLD5000 (diverse daily-life scenes) and Cowen17 (emotionally evocative scenes with constrained object categories). While decreased performance was observed for EmoNet fc8 compared to fc7 for the BOLD5000 dataset, *t*(14) = 25, *p* < 0.001, paired t-test (**Fig. 4e**), predictive performance increased for EmoNet fc8 compared to fc7 for the Cowen17 dataset, *t*(14) = 17.62, *p* < 0.001, paired t-test (**Fig. 4f**). These results indicate that object-level representations in fc7 of EmoNet—which were originally trained for object recognition— are critical for predicting emotion ratings in daily-life scenes of BOLD5000. However, the performance gain from fc7 to fc8 of EmoNet in Cowen17 suggests that abstract emotion-related categorical information encoded in fc8 provides additional predictive power beyond object-level features.

### Representational similarity of emotional information is greater within than between object categories

Having established that object categories (the fc8 layer in the AlexNet model) exhibit the strongest emotion predictions, we further investigated whether these features encode emotional information by organizing representations in a way that reflects emotional distinctions. We tested whether the representational similarity of emotional information within an object cluster is greater than that between object clusters. For instance, related object categories such as pizza and burger might exhibit higher emotional representational similarity compared to unrelated categories like pizza and car. Additionally, we sought to determine whether the fc8 layer encodes more emotional information compared to the early conv1 layer. If this is the case, the difference in pattern similarity values between within-cluster and between-cluster comparisons should be larger for the fc8 layer than for the conv1 layer. To explore these hypotheses, we applied k-means clustering across four different cluster numbers (k = 20, 30, 40, 50). The k-means algorithm was executed 10 times with varying initial cluster centroids and the best clustering result was chosen. Images within a cluster means similar images in terms of fc8 layer or conv1 layer. We computed representational similarities (Pearson correlations) between emotion ratings within and between clusters for both the conv1 and fc8 layers.

We observed significant differences between within-cluster and between-cluster similarities across various cluster numbers for both conv1 layer and for the fc8 layer (*p*s < 0.001, FDR corrected; **Fig. 5a-b**; **Supplementary text**). We then computed the difference between within-cluster and between-cluster similarities for each cluster number and conducted paired t-tests to compare these differences between the conv1 and fc8 layers. Significant differences were found between the fc8 and conv1 layers across various cluster numbers (*p*s < 0.001, FDR corrected; **Fig. 5c**; **Supplementary text**). These results demonstrated that the representational similarities of emotional information within object clusters were greater than those between object clusters, suggesting that related object categories share common emotional features.

**Fig. 5.**
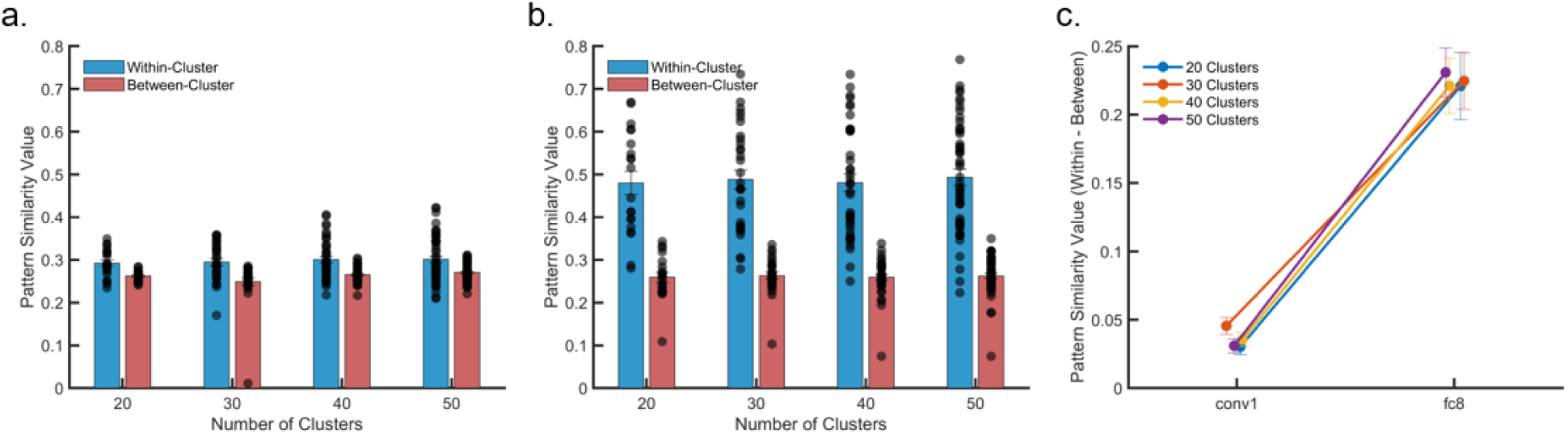
Representational similarities of emotional information within object clusters were greater than those between object clusters. The representational similarity analysis results indicated that both a) the conv1 layer and b) the fc8 layer of AlexNet exhibited greater within-cluster similarity than between-cluster similarity, suggesting that both layers encode emotional information, regardless of the number of K-means clusters. c) The fc8 layer demonstrated a larger difference in pattern similarity between within-cluster and between-cluster comparisons than the conv1 layer, indicating that the fc8 layer encodes more emotional information. Error bars represent the standard error across clusters and each dot represents the result of a cluster.

### Activity patterns in object-selective cortex predict emotion ratings

The previous analyses showed that object category representations reliably predict emotion ratings of daily-life scenes, suggesting that emotion recognition can rely on established object processing systems, at least for common, everyday scenes and objects. If emotion recognition relies primarily on the visual object recognition system, we would expect the object-selective visual cortex (i.e., the lateral occipital complex) to outperform all other visual regions (e.g., early visual cortex or scene-selective regions) in predicting emotional ratings.

To test this hypothesis, we analyzed the fMRI data from the BOLD5000 dataset, which includes data from four individuals who observed approximately 5000 unique images (Chang et al., 2019). The fMRI data comprise a complete dataset for three out of four participants across 15 functional sessions, with the fourth participant having data from 9 functional sessions. Each trial consisted of an image presented for 1 second, followed by a 9-second fixation cross. After viewing the image, participants were instructed to perform a valence judgment task, in which they rated their preference for the image by choosing “like,” “neutral,” or “dislike.” (**Fig. 6a-b**). The functional localizer sessions consisted of three types of conditions: scenes, objects, and scrambled images, with stimuli that were distinct from those used in the main fMRI study (**Fig. 6b**). The following regions of interest were included: Early visual cortex (EarlyVis); object selective lateral occipital complex (LOC); and scene selective regions of interest, including the parahippocampal place area (PPA), the retrosplenial complex (RSC), and the occipital place area (OPA). We included Heschl’s gyrus (Heschl) as a control region because it is a primary sensory area for the auditory modality, and it is not expected to significantly contribute to visual emotion representation (**Fig. 6c**).

**Fig. 6.**
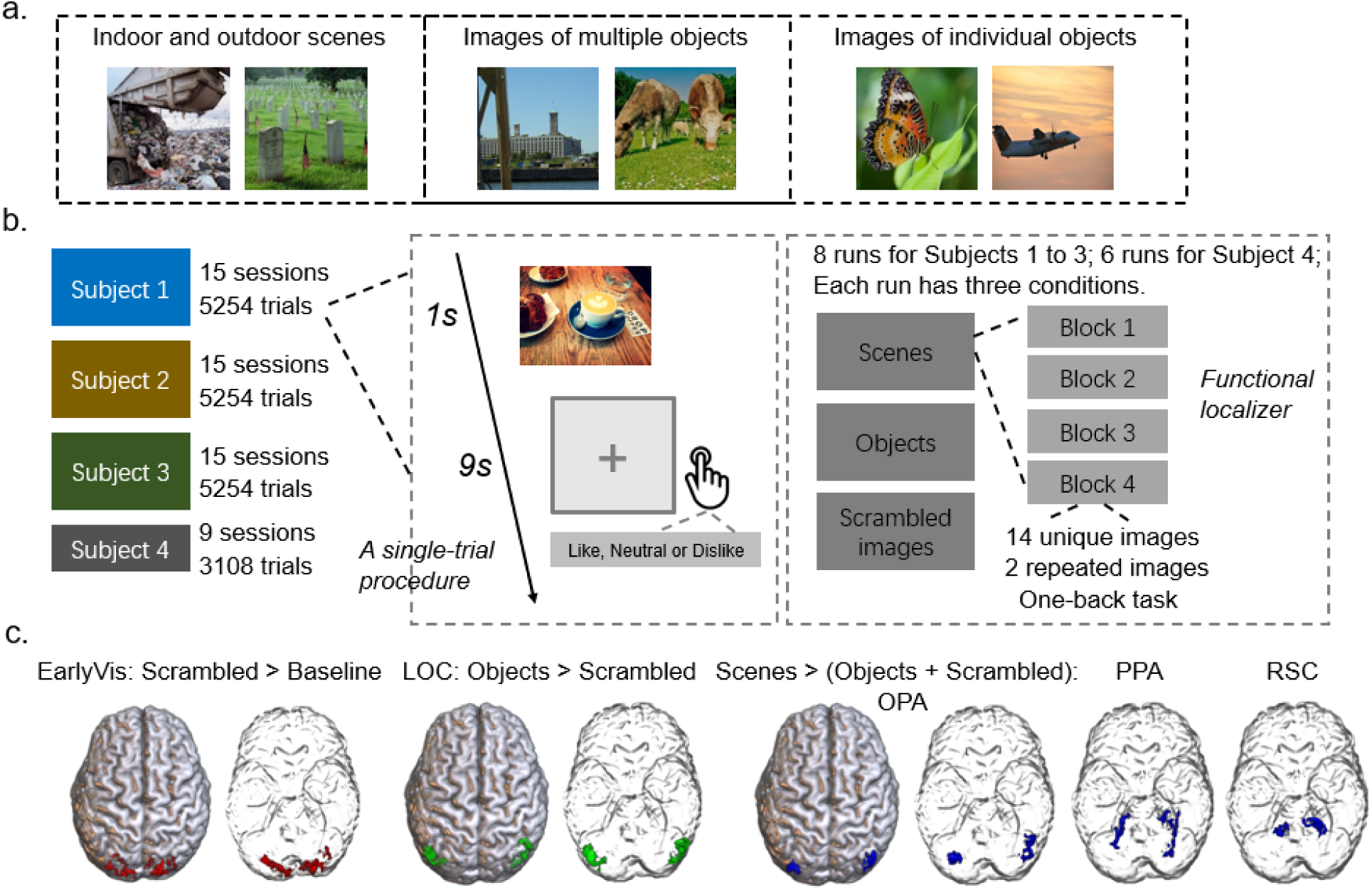
BOLD5000 experimental procedure and the regions of interest from the functional localizer. a) Experiment 1 stimuli consisted of 1000 images depicting indoor and outdoor scenes with a general focus rather than on specific objects, actions, or people; 2000 complex images featuring multiple objects, where these objects were often situated within a realistic context and depicted as interacting with other animate or inanimate entities; and 1916 images predominantly showcasing individual objects. b) The main fMRI data comprise a complete dataset for three out of four participants across 15 functional sessions, with the fourth participant having data from 9 functional sessions. Eight functional localizer sessions were conducted (six sessions for participant CSI4. c) The regions of interest from the functional localizer were defined using the following way. Early visual cortex was defined by comparing scrambled images to baseline. An object-selective regions of interest, the lateral occipital complex (LOC) was defined by comparing objects to scrambled images. Scene-selective regions of interest included the parahippocampal place area (PPA), the retrosplenial complex (RSC), and the occipital place area (OPA), defined by comparing scenes with objects and scrambled images.

We used PLSR to predict emotion ratings from activity patterns in the visual cortex (**Fig. 7a**). We found that all visual regions could significantly predict emotion ratings (**Fig. 7b-f**; **Supplementary text**; **Table S1**; **Fig. S6**). To compare the effects of region (Heschl, EarlyVis, LOC, OPA, PPA, and RSC), we performed repeated measures ANOVAs on the prediction-outcome correlation across different cross-validation folds, and Greenhouse-Geisser corrections were applied where necessary to account for violations of sphericity. We found main effects of region for all subjects: CSI1, *F*(4.61, 64.49) = 245.62, *p* < 0.001, partial = 0.95; CSI2, *F*(3.69, 51.61) =175.11, *p* < 0.001, partial = 0.93; CSI3, *F*(3.45, 48.27) = 258.91, *p* < 0.001, partial = 0.95; CSI4, *F*(2.61, 20.91) = 196.15, *p* < 0.001, partial = 0.96. These results indicate that different brain regions vary in their capability to predict emotion ratings.

**Fig. 7.**
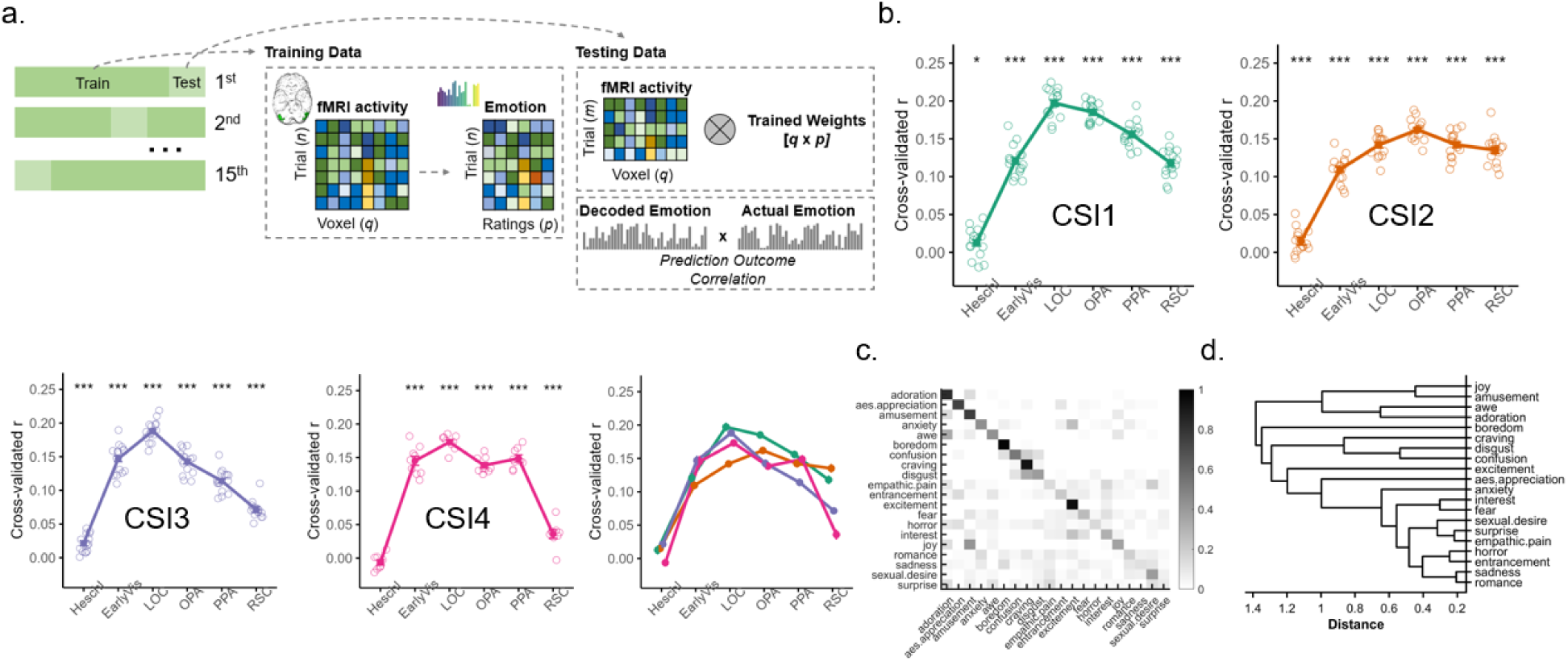
Object-selective cortex (LOC) outperformed other visual cortex regions in predicting emotion ratings. a) Decoding emotions from fMRI activity in visual cortices using partial least squares regression. b) Results of decoding emotions from fMRI activity in the visual cortices for CSI1, CSI2, CSI3, CSI4, and all subjects. fMRI activity in all visual cortex regions outperformed Heschl’s gyrus in predicting emotion ratings, and fMRI activity in the LOC (Lateral Occipital Complex) region outperformed that in the EarlyVis, OPA, PPA, and RSC regions in predicting emotion ratings. c) The averaged, normalized confusion matrix across four subjects for the relationship between the multivariate pattern responses in the LOC and emotion ratings. Rows represent the actual categories of the cross-validated data, and columns denote the predicted categories. Gray colormap indicates the proportion of predictions within the dataset, with each row summing to 1. Correct predictions fall on the diagonal of the matrix, whereas off-diagonal elements reflect wrong predictions. We correlated the predicted emotional ratings from the PLSR model for LOC cortical activity with vectors of emotional ratings across 20 emotion categories, resulting in a 20 × 20 correlation matrix. d) Dendrogram constructed using Ward’s method based on the confusion matrix in (c). *** p < 0.001; ** p < 0.01; * p < 0.05; ns, not significant.

Our main question was to investigate whether fMRI activity in the visual cortices provide better predictions of emotional ratings compared to activity in Heschl’s gyrus, and whether fMRI activity in the LOC region outperforms that in the EarlyVis, OPA, PPA, and RSC regions. We found that fMRI activity in all visual regions outperformed Heschl’s gyrus in predicting emotion ratings (**Fig. 7b-f**; **Supplementary text; Table S2**), and fMRI activity in the LOC region outperformed that in the EarlyVis, OPA, PPA, and RSC regions in predicting emotion ratings (**Fig. 7b-f**; **Supplementary text; Table S2**).

To further examine the representation of emotions within the LOC region, we constructed confusion matrices grounded in the PLSR model, using LOC cortical activity to predict emotion ratings. For each participant and each cross-validation iteration, we correlated the emotion ratings predicted by the PLSR model, based on LOC cortical activity, with a vector of actual emotional ratings spanning 20 distinct emotion categories. This process yielded a 20 × 20 correlation matrix per iteration. The emotion category with the highest correlation was designated as the most likely emotion inferred from brain activity. Subsequently, the confusion matrices derived from all cross-validation folds were aggregated. Finally, we normalized the confusion matrix to a scale between 0 and 1, and plotted the mean confusion matrix across all subjects (**Fig. 7g**). The data reveal several instances of emotion categorization that align with intuitive ambiguities, encompassing the conflation of joy with amusement, awe with adoration, anxiety with excitement, and interest with excitement. However, the findings also show several counterintuitive associations, notably the grouping of craving with disgust, and empathic pain with sexual desire (**Fig. 7g-h**). These findings suggest that while the LOC region does not perfectly capture human emotion ratings, it does contain a rich representation of emotion categories. Confusion matrices and dendrograms for other brain regions are presented in **Fig. S7**.

The fMRI results revealed that all visual regions significantly predicted emotion ratings, with object-selective regions outperforming other visual areas in their predictive ability. These findings are consistent with previous findings showing that successive layers in AlexNet lead to significant improvements in predicting emotion ratings. These results suggest that emotion ratings are driven by object representations encoded in the LOC region, indicating that emotion recognition primarily relies on the visual object recognition system.

## Discussion

Scenes and objects that we encounter in daily life are consistently associated with various emotions. However, the neural and computational mechanisms underlying the elicitation of diverse emotions by visual scenes remain unclear. To address this, we analyzed a large set of daily-life scenes along with a broad spectrum of object categories. We presented approximately 5000 images from this collection to 300 volunteers to obtain emotion ratings for each image. Analyzing this dataset allowed to evaluate whether emotions are evoked by an emotion-specific recognition system or whether they are driven by established mechanisms of object recognition.

We found that for explicitly emotionally evocative scenes from Cowen and Keltner (2017), EmoNet (an emotion-based visual system model) outperformed AlexNet (an object-based visual system model), replicating the findings of Kragel et al. (2019). However, AlexNet outperformed EmoNet in predicting emotion ratings for images of daily-life scenes and objects. This pattern was consistent across three subsets of the image dataset. Our findings thereby highlight the role of established object processing systems in driving emotion schemas for daily-life scenes.

If emotion recognition relies on established object processing systems for daily-life scenes, as our findings suggest, we would expect emotion information to be processed hierarchically, in line with the organization of the visual cortex for object recognition. Supporting this hypothesis, we observed a monotonic increase in the ability to predict emotion ratings with increasing layer depth in AlexNet, consistent with recent studies (Liu et al., 2024)^1^. While prior work has focused on broad valence categories (i.e., positive, neutral, and negative), we extended these findings (Liu et al., 2024) by demonstrating layered processing of emotional information using more fine-grained emotional categories. This layer-wise pattern was consistent across three distinct subsets of the BOLD5000 dataset, which varied in focus, with some subsets being more object-focused and others more scene-focused. Our representational similarity analyses further revealed that the emotion rating distances within object clusters, based on high-level object classes (from the fc8 layer of AlexNet), were greater than those between object clusters, indicating that object categories encode emotional information. The difference in pattern similarity between within-category and between-category comparisons was more pronounced in the fc8 layer than in the conv1 layer, indicating that the fc8 layer encodes more emotional information than earlier layers. These findings suggest that visual emotional representations of everyday scenes and objects are processed hierarchically in the visual object processing system, allowing for both rapid, coarse evaluations of emotional significance based on simple visual features (e.g., prototypical shapes and textures) and more detailed analyses of complex object features and concepts, depending on context (Öhman et al., 2001).

We found that EmoNet outperformed AlexNet in predicting emotion ratings for scenes from Cowen and Keltner (2017). These results suggest that object categories alone do not provide sufficient information for emotion prediction, although other visual features may play a critical role. When there is limited variability in the object categories present in the scenes (e.g., predominantly humans, as in the images used to train EmoNet), the final layer of AlexNet, which reflects object categories, becomes less important for predicting emotions. It is important to note that EmoNet was developed by keeping all layers of AlexNet fixed, except for the final fully connected layer, which was retrained. Consequently, the features from EmoNet’s earlier layers (conv1 to fc7) were originally trained for object categorization. Therefore, EmoNet performance still makes use of object representations, though not as abstract as those encoded in AlexNet’s final layer. Consistent with this, our results revealed that object-level representations in the fc7 layer of EmoNet/AlexNet are critical for predicting emotion ratings in daily-life scenes from BOLD5000. However, the fc8 layer of EmoNet outperformed the fc7 layer in predicting emotion ratings for the Cowen and Keltner (2017), suggesting that abstract emotion-specific categorical information encoded in fc8 provides additional predictive power beyond object-level features.

If emotion recognition primarily relies on the visual object recognition system, we would expect the object-selective visual cortex, specifically the LOC, to outperform other visual regions (e.g., early visual cortex) in predicting emotion ratings. To test this, we analyzed fMRI data from a comprehensive set of 5000 images alongside emotion ratings reported while viewing these images. Our results showed that fMRI activity in all visual regions, including EarlyVis, LOC, OPA, and PPA, outperformed Heschl’s gyrus (an auditory region) in predicting emotion ratings. These findings replicate previous research (Kragel et al., 2019), demonstrating that activity in the visual cortex significantly predicts emotion ratings for images (Abdel-Ghaffar et al., 2024; Bo et al., 2021). Our study provides novel insights by analyzing a larger and more diverse set of stimuli, capturing a broader range of variance across visual images, rather than relying on smaller sets of tens or hundreds of stimuli. This approach offers several unique benefits for uncovering universal principles of human brain function. Importantly, fMRI activity in the LOC exceeded that of other visual regions in predicting emotion ratings, suggesting that emotion ratings are primarily driven by object representations encoded in the LOC region (Epstein & Baker, 2019; Grill-Spector et al., 2001; Grill-Spector & Malach, 2004; Malach et al., 1995). Together with the demonstrated link between object categories and emotion and the findings of hierarchical processing of emotional information in AlexNet, these results collectively illustrate that emotion processing in everyday scenes and objects depends on the visual object recognition system.

These findings support theories emphasizing that object categorization is a necessary condition for emotional responses to visual scenes (Lazarus, 1982, 1984; Storbeck & Clore, 2007; Storbeck et al., 2006). They are in line with recent evidence showing that object recognition precedes the onset of affective feelings (Nummenmaa et al., 2010), with the two latencies being positively correlated across participants (Reisenzein & Franikowski, 2022). In addition, experimental manipulations that delayed object recognition also delayed the onset of affective feelings, and this effect was at least partially mediated by the delayed recognition of objects (Franikowski et al., 2021; Franikowski & Reisenzein, 2023). Furthermore, studies have shown that subjective valence and arousal ratings were modulated by the affective content of a scene only when the scene’s content was correctly reported; no affective modulation occurred when the picture content was not accurately identified (Mastria et al., 2024). A recent study also demonstrated that object categorization is necessary for the late positive potential (an emotion-related EEG marker) to be evoked (Codispoti et al., 2021). Our findings contribute to this literature by showing superior predictive performance of emotion ratings based on object content, supporting a strong link between object categories and emotion, even when using finer differentiation of emotional categories (over 20) compared to previous studies.

A potential limitation of the current study is the discrepancy between the fMRI task (valence rating) and the emotion categorization task, as task demands can modulate neural processing (Harel et al., 2014; Koc et al., 2025; VanRullen & Thorpe, 2001). The valence-rating task for the fMRI data diverges from both typical naturalistic perception in daily life and the emotion categorization task employed in our rating study. This divergence raises questions about the direct comparability of neural processes across tasks. However, our results indicate that despite these differences, visual cortical activity can predict emotion ratings successfully, suggesting that core emotion representations are preserved even when task demands shift. In addition, our results cannot exclusively be interpreted as supporting a causal relationship between object categorization (cause) and emotion (effect). It remains possible that object features facilitate emotion responses, or that affective responses assist in object recognition (Barrett & Bar, 2009). Future studies combining our approach with EEG/MEG could help to clarify the temporal stages at which these effects manifest.

## Method

### Behavioral Study

#### Participants

A total of 300 volunteers (143 females, 155 males, and 2 individuals who did not report their sex; mean age = 27.78 ± 4.38 years, and one participant who did not provide age information), recruited through Prolific, were included in the analyses. Additionally, six participants participated in the experiment but did not complete it due to technical issues or other reasons. These individuals were compensated but not included in the analyses. Participants were required to be over 18 years of age, possess normal or corrected-to-normal vision, and be fluent in English. This experiment received approval from the Ethics Committee of the Faculty of Social Sciences at Radboud University (ECSW-LT-2022-8-4-35788).

#### Stimuli

The stimuli included 4913 images, selected from the BOLD5000 public fMRI dataset involving human subjects, which contains 5254 images, with 4916 being unique (Chang et al., 2019). These stimuli featured real-world scenes, showcasing a broad diversity of images. They encompassed both outdoor and indoor scenes, complex interactions among objects, human social interactions, and objects situated within real-world contexts. Three out of the 4,916 unique images were not included in the experiment due to a technical issue. Additionally, to prepare participants for the main experiment, we introduced 10 new images from the OASIS affective image database (Kurdi et al., 2017) in a practice task.

#### Procedure

The 4913 images were randomly divided into 30 sets; 29 of these sets contained 165 images each, while one set comprised 128 images. Participants were recruited to evaluate each set individually, with a target of 10 participants per set. Additionally, for each set, 8 images were shown twice to assess the consistency of the ratings. At the beginning of the experiment, participants were briefed that they would be shown a series of images individually. Their task involved identifying the emotions each image elicited in them. After each image was displayed, participants were presented with a prompt featuring 20 emotion categories: adoration, aesthetic appreciation, amusement, anxiety, awe, boredom, confusion, craving, disgust, empathic pain, entrancement, excitement, fear, horror, interest, joy, romance, sadness, sexual desire, and surprise. These categories, chosen based on prior research, have been thoroughly validated as distinct by human evaluators (Kragel et al., 2019). Participants were instructed to select at least one category that best reflected their feelings towards the image, although they could select as many as desired. To ensure participants fully understood the meaning of each emotion category, they were guided through detailed descriptions of all 20 categories. Subsequently, they undertook 10 practice trials to become accustomed to the procedure.

The main part of the experiment was divided into eight blocks. The first seven blocks each contained 20 images, with the remainder displayed in the final block. There was a self-paced rest period between each block. Every trial began with a 250 ms fixation cross, succeeded by the presentation of an image for one second. This was followed by the emotion category prompt. If participants were uncertain about the meaning of any emotion while responding, they could view a description of each emotion by moving the mouse over the emotion category labels. Afterwards, a prompt would inquire about their confidence in their selection. Both the emotion categorization and confidence assessment were conducted at the participants’ own pace. The emotional ratings data are available at Open Science Framework (https://osf.io/eks8u/?view_only=b51e2d80b41748329ad9009343438c6d).

### fMRI Dataset

#### Participants

The fMRI data were sourced from the BOLD5000 dataset (Chang et al., 2019), which are publicly available at: https://kilthub.cmu.edu/articles/dataset/BOLD5000_Release_2_0/14456124Participants were graduate students from Carnegie Mellon University. The demographic details of the participants are as follows: CSI1, a 27-year-old male; CSI2, a 26-year-old female; CSI3, a 24-year-old female; and CSI4, a 25-year-old female. All participants were right-handed, with no reported history of psychiatric or neurological disorders, nor any current psychoactive medication use. Each participant provided written informed consent, adhering to protocols approved by the Institutional Review Board at Carnegie Mellon University.

#### Stimuli

The stimuli consisted of 4916 unique images, categorized as follows: 1000 hand-selected images depicting indoor (for example, restaurants) and outdoor (such as mountains and rivers) scenes with a general focus rather than on specific objects, actions, or people; 2000 complex images featuring multiple objects, where these objects were often situated within a realistic context and depicted as interacting with other animate or inanimate entities (for instance, scenes of human social interactions); and 1916 images predominantly showcasing individual objects. The luminance of all images was standardized through a gray-world normalization technique.

#### Procedure

##### Main fMRI experiment

The fMRI data include a full set of recordings for three of the four participants across 15 functional sessions, while the fourth participant’s data cover 9 functional sessions.

These sessions are divided into two types: eight sessions each consisting of 9 runs, and seven sessions comprising 10 runs each. Every run included 37 trials, with each trial featuring an image displayed for 1 second followed by a 9-second fixation cross. After the stimulus presentation, participants were asked to perform a valence judgment task, where they evaluated their preference for the image by selecting “like,” “neutral,” or “dislike.” This was accomplished using an MRI-compatible response glove on their dominant hand.

##### Functional localizer

Eight functional localizer sessions were conducted (six sessions for participant CSI4). These localizers were conducted at the end of the day’s session when a participant had completed a main functional session of 9 runs, but not those with 10 runs. The functional localizer sessions included three types of conditions: scenes, objects, and scrambled images, and the stimuli presented did not overlap with the images used in the main fMRI study. There were four blocks per condition, with each block containing 16 trials that included 14 unique images and 2 repeated images (**Fig. 1b**). Participants were asked to perform a one-back task, which required them to press a button if an image was immediately repeated.

##### MRI acquisition

MRI data were collected using a 3T Siemens Verio MR scanner at Carnegie Mellon University. Functional images were acquired through a T2*-weighted gradient-recalled echo-planar imaging multi-band pulse sequence. The in-plane resolution was set at 2 x 2 mm, with a matrix size of 106 x 106. The slice thickness was 2 mm without any gap. The field of view was 212 mm. The repetition time (TR) was 2000 ms, echo time (TE) was 30 ms, and the flip angle was set at 79 degrees. The multi-band factor was 3.

Additional details can be found in the original paper (Chang et al., 2019).

##### fMRI data preprocessing

We analyzed Release 2.0 of BOLD5000 (version descriptor: TYPED-FITHRF-GLMDENOISE-RR), which employed GLMsingle, a denoising toolbox (Prince et al., 2022). This toolbox includes a custom hemodynamic response function estimation, GLM denoising (Kay et al., 2013), and regularization via ridge regression. It has been shown that this approach greatly enhances the quality of data. Before the GLMsingle denoising procedure was applied, the data were preprocessed using fMRIPrep (Esteban et al., 2019).

Functional localizer data were also preprocessed using fMRIPrep (Esteban et al., 2019). Subsequently, SPM12 was employed for GLM analyses, incorporating three conditions: scenes, objects, and scrambled images (Penny et al., 2011). Additionally, nine nuisance regressors were included, comprising six motion parameters, the average signal within the cerebral spinal fluid mask and white matter mask, as well as global signal within the whole-brain mask.

### Data analyses

#### Emotional rating analyses

##### Computation of emotion probabilities

For each image, we compiled the frequency of selected emotion categories from 10 participants. The probability for each of the 20 emotion categories was calculated by dividing the number of times an emotion category was selected by the total number of participants, which is 10. For instance, for a particular image, if ‘joy’ was selected by 4 out of 10 participants, ‘adoration’ by all 10, and ‘disgust’ by none, the probability for ‘joy’ would be 40%, for ‘adoration’ 100%, and for ‘disgust’ 0%. This approach establishes the maximum probability of any emotion evoked by an image at 100% and the minimum at 0% without assuming that emotion categories are mutually inclusive or exclusive.

##### t-Distributed stochastic neighbor embedding (t-SNE)

To explore the structure of images, we used the Barnes-Hut version of t-SNE, setting the perplexity to 30 and theta to 0, as an approach to visualize data in high dimensions. This approach took a normalized matrix of 4913 observations by 20 emotions of emotion probabilities and computed pairwise Euclidean distances between observations. These Euclidean distances in high-dimensional space between variables were then transformed into conditional probabilities. These probabilities are then mapped onto a two-dimensional space through the Student’s t-distribution, aiming to minimize the Kullback–Leibler divergence in the process. This method has the advantage of revealing the global structure but also capturing the local structure within the high-dimensional data (Van der Maaten & Hinton, 2008).

##### Emotion dissimilarity

To assess how different categories of emotions are related, we used the matrix of emotion probabilities of 4913 images by 20 emotions as input and calculated the pairwise Pearson correlation distance between two emotions across all evaluated images. We chose the Pearson correlation distance for its direct relationship with the squared Euclidean distances of normalized vectors, which simplifies understanding. To calculate dissimilarity, we subtracted the Pearson correlation coefficient from 1.

##### Hierarchical clustering analysis

To further visualize how emotion categories cluster together based on the probabilities associated with the 20 distinct emotion categories, we conducted an analysis of hierarchical clustering. This analysis was carried out utilizing Ward’s method as the criterion (Murtagh & Legendre, 2014).

### Comparing object and emotion deep convolutional neural network (DCNN) representations in predicting emotions

To test whether emotion recognition relies on emotion-specific visual processing or can similarly be explained by established object and scene processing systems, we compared the performance of object and emotion DCNN representations in predicting emotions. We used the EmoNet DCNN model as an emotion deep convolutional neural network model to extract features from 4913 images. EmoNet is a convolutional neural network derived from AlexNet (Krizhevsky et al., 2012, 2017), with its objective modified from object class recognition to the classification of images into 20 distinct emotion categories. This was achieved by retraining the weights in its final fully connected layer (Kragel et al., 2019). The AlexNet DCNN model was implemented via MATLAB’s Deep Learning Toolbox, to extract features from 4913 images. The number of units in each layer of AlexNet is: conv1, 96x55x55; conv2, 256x27x27; conv3 and conv4, 384x13x13; conv5, 256x13x13; fc6 and fc7, 4096; and fc8, 1000. For convolutional layers, we averaged activations over the spatial domain, resulting in vector lengths of 96, 256, 384, 384, and 256, respectively. This averaging reduces dimensionality, aligns convolutional outputs with the fully connected layers, and emphasizes the global presence of features over their exact spatial locations (see Lindh et al., 2019, for a similar approach).

Using partial least squares regression (PLSR), we predicted emotion probabilities from features extracted from the fc8 layer of EmoNet or AlexNet in a leave-one-session-out cross-validation approach. PLSR analyses were conducted using the ‘plsregress’ function in MATLAB. The prediction performance was assessed by calculating the Pearson correlation between observed and cross-validated predicted emotion ratings, with permutation testing employed for inference.

### Comparative analyses across three subsets of BOLD5000 images

To evaluate the consistency of the comparison between AlexNet and Emonet across diverse image datasets, we conducted a comparative analysis using three distinct subsets of BOLD5000 images: Scenes (Xiao et al., 2010), COCO (Lin et al., 2014), and ImageNet (Russakovsky et al., 2015). Specifically, we conducted two analyses: first, decoding emotions from AlexNet fc8 representations, and second, decoding emotions from the fc8 layer in the EmoNet model. The analysis procedure was identical to the previous analyses, with the exception that 5-fold cross-validation was used instead of 15-fold due to the smaller number of images in each of the three subsets.

In addition, we analyzed videos from Cowen and Keltner (2017), following Kragel et al. (2019). The 2185 videos, along with their mean emotion ratings from 853 participants, were sourced from Cowen and Keltner (2017). To be consistent with our other analyses and the previous study (Kragel et al., 2019), we selected 20 emotion categories. We extracted three representative frames from each video at 25%, 50%, and 75% of the video length, resulting in 6555 images. We then performed the two analyses described above on these 6555 images. The analysis procedure was consistent with the previous analyses, employing a 15-fold cross-validation method, where the dataset was partitioned into 15 folds.

### Decoding emotions from AlexNet layer activations

To determine whether emotional information is processed in a hierarchical manner, we predicted emotion probabilities based on activations from various layers of AlexNet. We implemented PLSR models with 20 dimensions to prevent overfitting, in accordance with previous research (Čeko et al., 2022; Soderberg et al., 2023). PLSR models were trained using a leave-one-session-out cross-validation approach. PLSR was employed to establish a linear relationship between layer activations and emotion ratings within the training set. This transformation was then applied to the activations from the test set (the left-out session) to decode emotion probabilities for each image in that session. To evaluate out-of-sample prediction performance, we calculated the Pearson correlation between observed and cross-validated predicted emotion ratings. Inference on prediction performance was conducted using permutation testing, involving 1000 iterations of PLSR analyses with random shuffling of emotion ratings to generate a null distribution of correlation coefficients (r values). Statistical significance was determined by comparing the actual r values to this permutation distribution, and multiple comparisons were corrected using the false discovery rate (FDR) method (Benjamini & Yekutieli, 2001).

### The link between object deep neural network representations and emotional ratings via representational similarity analyses

To investigate whether object features encode emotional information by organizing representations that reflect emotional distinctions, we evaluated whether the representational similarity of emotional information within an object cluster exceeds that between object clusters. To achieve this, we first performed k-means clustering analyses to ensure a larger number of images within each object cluster. We used multiple replicates (10 in this case) to minimize the risk of converging on a suboptimal clustering solution due to an unfavorable initial configuration. The selected cluster numbers were chosen to cover a range of potential cluster granularities, from relatively coarse (k = 20) to finer (k = 50) groupings. This approach allowed us to assess how clustering structure and the resulting emotion similarity metrics might vary with the number of clusters.

We calculated representational similarities (using Pearson correlations) between emotion ratings both within and across clusters for the conv1 and fc8 layers. Within-cluster similarity was calculated as the average of the lower triangle correlations in the Pearson correlation matrix for stimuli within the same cluster, while between-cluster similarity was the average correlation for stimuli across different clusters. Paired t-tests were conducted to evaluate the significance of within- versus between-cluster similarities for each layer individually. Furthermore, we computed the difference between within- and between-cluster similarities for each cluster number and performed paired t-tests to compare these differences between the conv1 and fc8 layers. All p-values were corrected for multiple comparisons using the FDR method (Benjamini & Yekutieli, 2001).

### Regions of interest

The regions of interest from the functional localizer were defined using the following way. Early visual cortex (EarlyVis) was defined by comparing scrambled images to baseline. Lateral occipital complex (LOC) was defined by comparing objects to scrambled images. Parahippocampal place area (PPA), the retrosplenial complex (RSC), and the occipital place area (OPA) were defined by comparing scenes with objects and scrambled images (See Chang et al., 2019 for more details). Heschl’s gyrus was identified using the Anatomical Automatic Labeling (AAL) system version 3 (www.gin.cnrs.fr/en/tools/aal/).

### Decoding emotions from fMRI activity in visual cortices

Before conducting PLSR analyses, both the predictors (voxel-wise fMRI activity) and emotion probabilities (emotion probabilities on 20 dimensions) were normalized across trials for each participant in the fMRI experiment to yield a mean of 0 and a standard deviation of 1. PLSR analyses were then conducted for each participant (CSI1, CSI2, CSI3, and CSI4) and each ROI (Heschl, EarlyVis, LOC, OPA, PPA, and RSC) in a leave-one-session-out cross-validated approach. It is worth noting that CSI4 had a different number of sessions compared to the other three participants, resulting in a varied number of folds for cross-validation: 15-fold cross-validation for CSI1, CSI2, and CSI3, and 9-fold cross-validation for CSI4.

PLSR was used to establish a linear transformation between fMRI components and emotion components within a training set. This learned transformation was then applied to the fMRI components from the test set (i.e., the left-out session) to decode emotion probabilities for each image within that session.

To assess out-of-sample prediction performance, we computed the Pearson correlation between observed and cross-validated predicted emotion ratings. Inference on prediction performance was made using permutation testing. This involved rerunning the PLSR analyses using the same data and analytic procedure, but with random shuffling of the emotion ratings. This process was iterated 1000 times to generate a null distribution of correlation coefficients (r values). Statistical significance was determined by comparing the true r values to the permutation distribution. The false discovery rate (FDR) was used to correct for multiple comparisons (Benjamini & Yekutieli, 2001).

## Footnotes

1 It should be noted that AlexNet does not perfectly align with the human visual system due to its relatively few layers, limited recurrent processing, and other architectural differences.

## Notes

### Competing Interest Statement

The authors have declared no competing interest.

### Summary of Updates

Figure 4 revised; Manuscript text updated; Supplemental files updated.

