## Supplemental information for "Object representations drive emotion schemas across a large and diverse set of daily-life scenes"

**This file includes:**

*Supplementary text:*

*Individual and joint probability of emotion ratings;*

*Dissimilarity matrix and hierarchical clustering results;*

*Monte Carlo simulation for analyses of concordance in judgment rates;*

*The predictive performances of different layers in AlexNet for emotion ratings;*

*Results of layered processing of emotion information in AlexNet;*

*Results of representational similarity analyses of emotion information for object categories;*

*Results of predicting emotion ratings from visual cortical activity.*

*Table S1.*

*Table S2.*

*Figure S1.*

*Figure S2.*

*Figure S3.*

*Figure S4.*

*Figure S5.*

*Figure S6.*

**Supplementary text:**

**Individual and joint probability of emotion ratings**

The mean probability with which each emotion category was chosen over 4913 images was shown in **Fig. S1a**. As shown in **Fig. S1a**, "interest" and "boredom" were the most frequently selected labels, followed by emotions such as "joy," "aesthetic appreciation," and "adoration." To quantify relationships between emotion pairs, we calculated their joint probability (**Fig. S1b**). For each image, the joint probability of emotions A and B was derived by multiplying their individual probabilities, then averaging these values across all 4913 images. We subtracted the joint probabilities from 1 to obtain the complement of the joint probability. However, this method may overemphasize emotions with inherently high individual probabilities or outliers in specific images, even if their selection patterns are uncorrelated. Emotion dissimilarity can otherwise quantify the consistency with which pairs of emotions are selected across images. This analysis enables a more accurate evaluation of the relationships between distinct emotion categories. Low dissimilarity indicates emotions share similar selection patterns (e.g., both frequently chosen in certain image types and rarely in others).

**Dissimilarity matrix and hierarchical clustering results**

To assess how different categories of emotions are related, we evaluated the dissimilarity between each pair of emotions. The emotion dissimilarity matrix showed meaningful relationships between different emotion categories. For example, ‘Joy’, ‘Amusement’, ‘Adoration’ and ‘Interest’ were related to each other (**Fig. S2a**). To further visualize how emotion categories cluster together based on the probabilities associated with the 20 distinct emotion categories, we conducted an analysis of hierarchical clustering. Hierarchical clustering analyses showed meaningful clustering of emotion categories. For example, ‘Horror’, ‘Fear’, and ‘Anxiety’ were clustered together (**Fig. S2b**).

**Monte Carlo simulation for analyses of concordance in judgment rates**

We evaluated the reliability of emotion ratings for images by assessing the proportion of pictures exhibiting significant concordance in judgment rates across the 20 emotion categories following Cowen et al. (2017). Concordance here refers to the agreement among multiple raters in assigning a given image to the same emotion category. To determine the significance of the judgment proportions, we generated a null distribution of judgements via Monte Carlo simulation. Specifically, we simulated random 20-category judgments of 100,000 picture samples from a multinomial distribution with the probabilities for each emotion category to the actual proportion of times the category was selected for our 4913 images. We calculated the p-value as the proportion of simulated trials where the observed category proportion exceeded the null distribution.

**The predictive performances of different layers in AlexNet for emotion ratings**

The predictive performances of different layers in AlexNet for emotion ratings was as follows: conv1 (average leave-one-session-out cross-validated r = 0.121, p < 0.001, permutation test, FDR corrected), conv2 (average leave-one-session-out cross-validated r = 0.237, p < 0.001, permutation test, FDR corrected), conv3 (average leave-one-session-out cross-validated r = 0.297, p < 0.001, permutation test, FDR corrected), conv4 (average leave-one-session-out cross-validated r = 0.325, p < 0.001, permutation test, FDR corrected), conv5 (average leave-one-session-out cross-validated r = 0.339, p < 0.001, permutation test, FDR corrected), fc6 (average leave-one-session-out cross-validated r = 0.362, p < 0.001, permutation test, FDR corrected) , fc7 (average leave-one-session-out cross-validated r = 0.380, p < 0.001, permutation test, FDR corrected), and fc8 (average leave-one-session-out cross-validated r = 0.407, p < 0.001, permutation test, FDR corrected). Null distributions of correlation coefficients derived from permutations were presented in **Fig. S4**).

**Results of layered processing of emotion information in AlexNet**

We found that conv2 outperformed conv1 ($\Delta r$ = 0.116, *p* < 0.001, permutation test, FDR corrected), conv3 outperformed conv2 ($\Delta r$ = 0.060, *p* < 0.001, permutation test, FDR corrected), conv4 outperformed conv3 ($\Delta r$ = 0.028, *p* < 0.001, permutation test, FDR corrected), fc6 outperformed conv5 ($\Delta r$ = 0.023, *p* < 0.001, permutation test, FDR corrected), fc7 outperformed fc6 ($\Delta r$ = 0.018, *p* = 0.009, permutation test, FDR corrected), and fc8 outperformed fc7 ($\Delta r$ = 0.027, *p* < 0.001, permutation test, FDR corrected). No significant difference was found between conv5 and conv4 ($\Delta r$ = 0.014, *p* = 0.057, permutation test, FDR corrected).

**Results of representational similarity analyses of emotion information for object categories**

For the conv1 layer, we observed significant differences between within-cluster and between-cluster similarities across various cluster numbers: *t*(19) = 5.307, *p* < 0.001 (FDR corrected) when the cluster number was 20; *t*(29) = 7.349, *p* < 0.001 (FDR corrected) when the cluster number was 30; *t*(39) = 6.913, *p* < 0.001 (FDR corrected) when the cluster number was 40; and *t*(49) = 6.053, *p* < 0.001 (FDR corrected) when the cluster number was 50. Similarly, for the fc8 layer, significant differences between within-cluster and between-cluster similarities were also found: *t*(19) = 9.024, *p* < 0.001 (FDR corrected) when the cluster number was 20; *t*(29) = 10.872, *p* < 0.001 (FDR corrected) when the cluster number was 30; *t*(39) = 10.972, *p* < 0.001 (FDR corrected) when the cluster number was 40; and *t*(49) = 12.890, *p* < 0.001 (FDR corrected) when the cluster number was 50.

We then computed the difference between within-cluster and between-cluster similarities for each cluster number and conducted paired t-tests to compare these differences between the conv1 and fc8 layers. Significant differences were found between the fc8 and conv1 layers across various cluster numbers: *t*(19) = 7.694, *p* < 0.001 (FDR corrected) when the cluster number was 20; *t*(29) = 8.663, *p* < 0.001 (FDR corrected) when the cluster number was 30; *t*(39) = 9.308, *p* < 0.001 (FDR corrected) when the cluster number was 40; and *t*(49) = 10.447, *p* < 0.001 (FDR corrected) when the cluster number was 50. These results indicate that the representational distances of emotion information within object clusters were greater than those between object clusters, and that the deeper layer (i.e., fc8) encoded more emotion information than the earlier layer (i.e., conv1).

**Results of predicting emotion ratings from visual cortical activity**

First, we found that all visual regions can significantly predict emotion ratings, and comparable results were found for CSI1, CSI2, CSI3 and CSI4 (**Table S1**; Null distributions of correlation coefficients derived from permutations were presented in **Fig. S5**).

In addition, our study aimed to investigate whether fMRI activity in the visual cortices provides better predictions of emotion ratings compared to activity in Heschl's gyrus, and whether fMRI activity in the LOC region outperforms that in the EarlyVis, OPA, PPA, and RSC regions. We found that fMRI activity in all visual regions including EarlyVis ($\Delta r$ = 0.108, *p* < 0.001, permutation test, FDR corrected), LOC ($\Delta r$ = 0.184, *p* < 0.001, permutation test, FDR corrected), OPA ($\Delta r$ = 0.172, *p* < 0.001, permutation test, FDR corrected), PPA ($\Delta r$ = 0.143, *p* < 0.001, permutation test, FDR corrected), and RSC ($\Delta r$ = 0.105, *p* < 0.001, permutation test, FDR corrected), outperformed Heschl’s gyrus in predicting emotion ratings for CSI1. We also found that fMRI activity in the LOC region outperformed that in EarlyVis ($\Delta r$ = 0.076, *p* < 0.001, permutation test, FDR corrected), OPA ($\Delta r$ = 0.012, *p* = 0.099, permutation test, FDR corrected; *p* < 0.05, uncorrected), PPA ($\Delta r$ = 0.041, *p* < 0.001, permutation test, FDR corrected), and RSC ($\Delta r$ = 0.079, *p* < 0.001, permutation test, FDR corrected), in predicting emotion ratings for CSI1. Comparable results were found for CSI2, CSI3 and CSI4 (**Table S2**). These results suggest that fMRI activity in the visual cortices predicted emotion ratings more effectively than activity in Heschl's gyrus. Additionally, fMRI activity in the LOC region outperformed that in the EarlyVis, OPA, PPA, and RSC regions in predicting emotion ratings.

***Table S1. Results of decoding emotions from fMRI activity in the visual cortices.***

| **Subject** | **Cross-validated *r*** | ***SE*** | ***p* value** |
| --- | --- | --- | --- |
| **CSI1** |  |  |  |
| *Heschl* | 0.013 | 0.004 | 0.012 |
| *EarlyVis* | 0.121 | 0.004 | < 0.001 |
| *LOC* | 0.197 | 0.004 | < 0.001 |
| *OPA* | 0.185 | 0.004 | < 0.001 |
| *PPA* | 0.156 | 0.004 | < 0.001 |
| *RSC* | 0.118 | 0.005 | < 0.001 |
| **CSI2** |  |  |  |
| *Heschl* | 0.015 | 0.005 | < 0.001 |
| *EarlyVis* | 0.110 | 0.003 | < 0.001 |
| *LOC* | 0.142 | 0.004 | < 0.001 |
| *OPA* | 0.162 | 0.003 | < 0.001 |
| *PPA* | 0.142 | 0.004 | < 0.001 |
| *RSC* | 0.135 | 0.005 | < 0.001 |
| **CSI3** |  |  |  |
| *Heschl* | 0.021 | 0.004 | < 0.001 |
| *EarlyVis* | 0.147 | 0.004 | < 0.001 |
| *LOC* | 0.188 | 0.003 | < 0.001 |
| *OPA* | 0.143 | 0.004 | < 0.001 |
| *PPA* | 0.114 | 0.003 | < 0.001 |
| *RSC* | 0.071 | 0.004 | < 0.001 |
| **CSI4** |  |  |  |
| *Heschl* | -0.006 | 0.004 | 0.863 |
| *EarlyVis* | 0.145 | 0.007 | < 0.001 |
| *LOC* | 0.173 | 0.003 | < 0.001 |
| *OPA* | 0.138 | 0.004 | < 0.001 |
| *PPA* | 0.149 | 0.005 | < 0.001 |
| *RSC* | 0.036 | 0.007 | < 0.001 |

Notes. SE indicates the standard error across different cross-validated folds.

***Table S2. Results of the comparative analysis of different brain regions in predicting emotion experiences.***

| **Subject** | $\boldsymbol{\Delta}\boldsymbol{r}$ | ***p* value** | **FDR-corrected *p* value** |
| --- | --- | --- | --- |
| **CSI1** |  |  |  |
| *EarlyVis > Heschl* | 0.108 | < 0.001 | < 0.001 |
| *LOC > Heschl* | 0.184 | < 0.001 | < 0.001 |
| *OPA > Heschl* | 0.172 | < 0.001 | < 0.001 |
| *PPA > Heschl* | 0.143 | < 0.001 | < 0.001 |
| *RSC > Heschl* | 0.105 | < 0.001 | < 0.001 |
| *LOC > EarlyVis* | 0.076 | < 0.001 | < 0.001 |
| *LOC > OPA* | 0.012 | 0.035 | 0.099 |
| *LOC > PPA* | 0.041 | < 0.001 | < 0.001 |
| *LOC > RSC* | 0.079 | < 0.001 | < 0.001 |
| **CSI2** |  |  |  |
| *EarlyVis > Heschl* | 0.095 | < 0.001 | < 0.001 |
| *LOC > Heschl* | 0.127 | < 0.001 | < 0.001 |
| *OPA > Heschl* | 0.147 | < 0.001 | < 0.001 |
| *PPA > Heschl* | 0.127 | < 0.001 | < 0.001 |
| *RSC > Heschl* | 0.120 | < 0.001 | < 0.001 |
| *LOC > EarlyVis* | 0.032 | < 0.001 | < 0.001 |
| *LOC > OPA* | -0.020 | > 0.99 | > 0.99 |
| *LOC > PPA* | 0.000 | 0.509 | > 0.99 |
| *LOC > RSC* | 0.007 | 0.146 | 0.531 |
| **CSI3** |  |  |  |
| *EarlyVis > Heschl* | 0.126 | < 0.001 | < 0.001 |
| *LOC > Heschl* | 0.167 | < 0.001 | < 0.001 |
| *OPA > Heschl* | 0.121 | < 0.001 | < 0.001 |
| *PPA > Heschl* | 0.093 | < 0.001 | < 0.001 |
| *RSC > Heschl* | 0.050 | < 0.001 | < 0.001 |
| *LOC > EarlyVis* | 0.041 | < 0.001 | < 0.001 |
| *LOC > OPA* | 0.046 | < 0.001 | < 0.001 |
| *LOC > PPA* | 0.074 | < 0.001 | < 0.001 |
| *LOC > RSC* | 0.117 | < 0.001 | < 0.001 |
| **CSI4** |  |  |  |
| *EarlyVis > Heschl* | 0.151 | < 0.001 | < 0.001 |
| *LOC > Heschl* | 0.179 | < 0.001 | < 0.001 |
| *OPA > Heschl* | 0.145 | < 0.001 | < 0.001 |
| *PPA > Heschl* | 0.155 | < 0.001 | < 0.001 |
| *RSC > Heschl* | 0.042 | < 0.001 | < 0.001 |
| *LOC > EarlyVis* | 0.028 | < 0.001 | < 0.001 |
| *LOC > OPA* | 0.035 | < 0.001 | < 0.001 |
| *LOC > PPA* | 0.024 | 0.001 | 0.003 |
| *LOC > RSC* | 0.137 | < 0.001 | < 0.001 |

Notes. FDR correction was applied over all 9 comparisons.

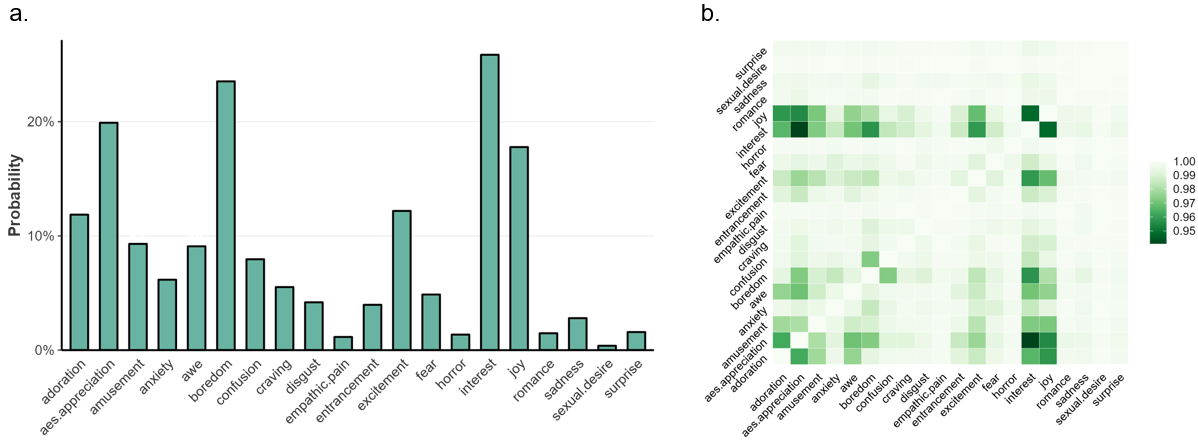

***Fig. S1. Individual and joint probability of emotion ratings.*** *a) Individual probability of emotion ratings. b) Joint probability of emotion ratings. We subtracted the joint probabilities from 1 to obtain the complement of the joint probability.*

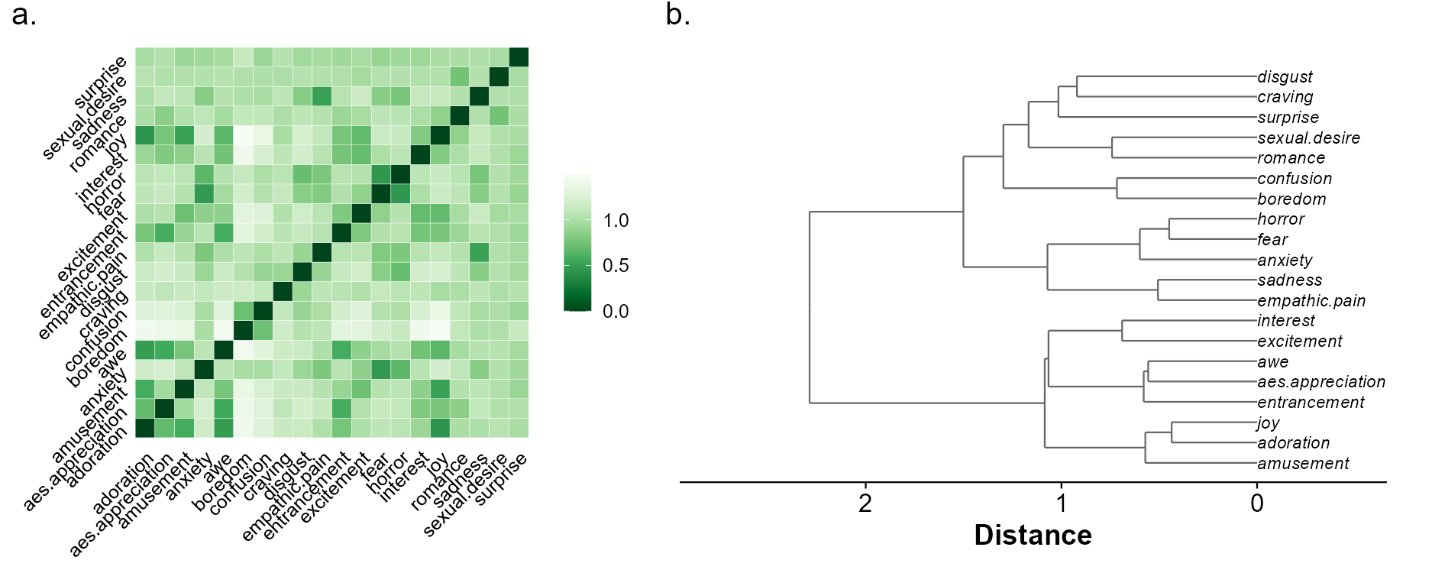

***Fig. S2. Dissimilarity matrix and hierarchical clustering results.*** *a) Emotion dissimilarity matrix, which was calculated using the Pearson correlation distance between two vectors that represent the probability of each emotion across all evaluated images. b) Hierarchical clustering results showed the clustering of emotion categories.*

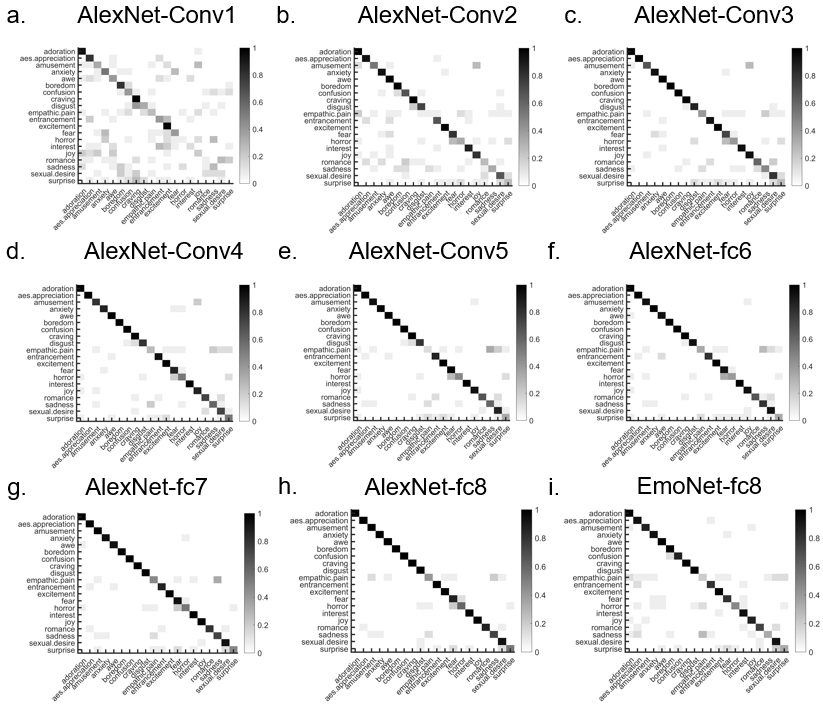

***Fig. S3. The averaged, normalized confusion matrices for decoding emotions from AlexNet and EmoNet layer activations.*** *Rows represent the actual categories of the cross-validated data, and columns denote the predicted categories. Gray colormap indicates the proportion of predictions within the dataset, with each row summing to 1. Correct predictions fall on the diagonal of the matrix, whereas off-diagonal elements reflect wrong predictions.*

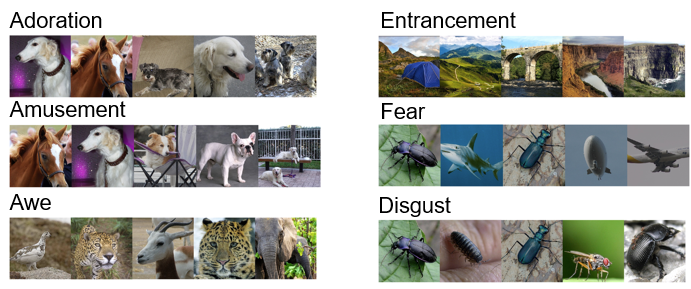

***Fig. S4. Top 5 example images predicted from the fc8 layer of AlexNet for 6 example emotions.***

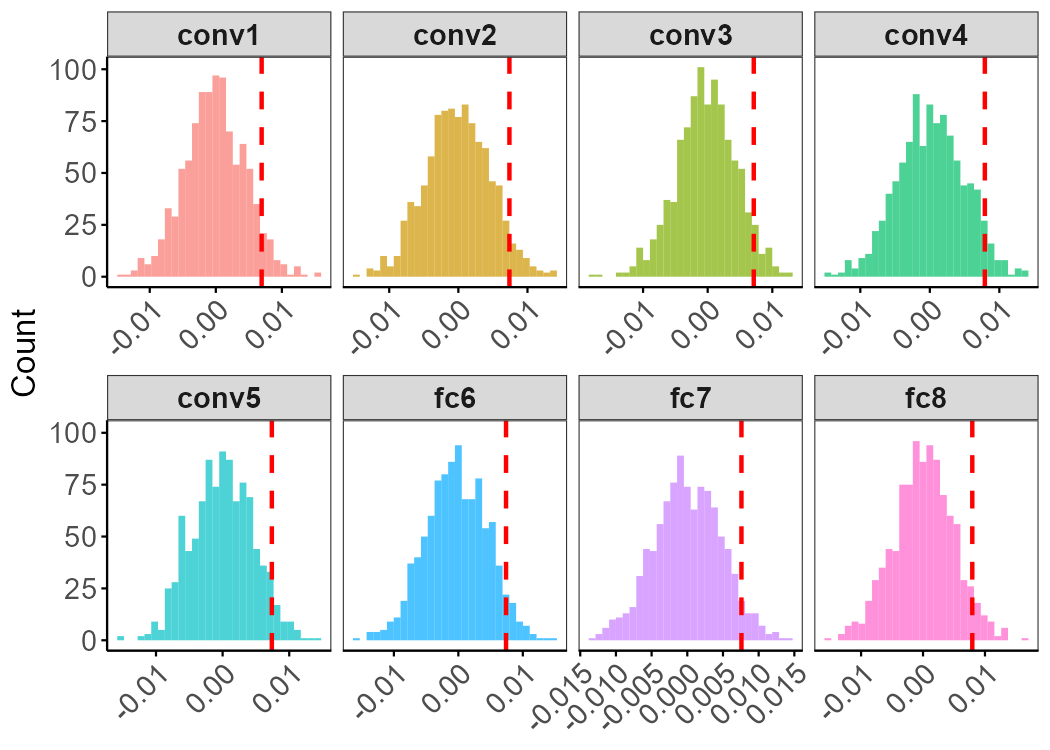

***Fig. S5. Null distributions of correlation coefficients derived from permutations for decoding emotions from object deep convolutional neural network (DCNN) representations.*** *This involved rerunning the Partial Least Squares Regression (PLSR) analyses using the same data and analytic procedure, but with random shuffling of the emotion ratings. This process was iterated 1000 times to generate a null distribution of correlation coefficients (r values). The red line on the x-axis of each subplot indicates the r values where q = 0.05.*

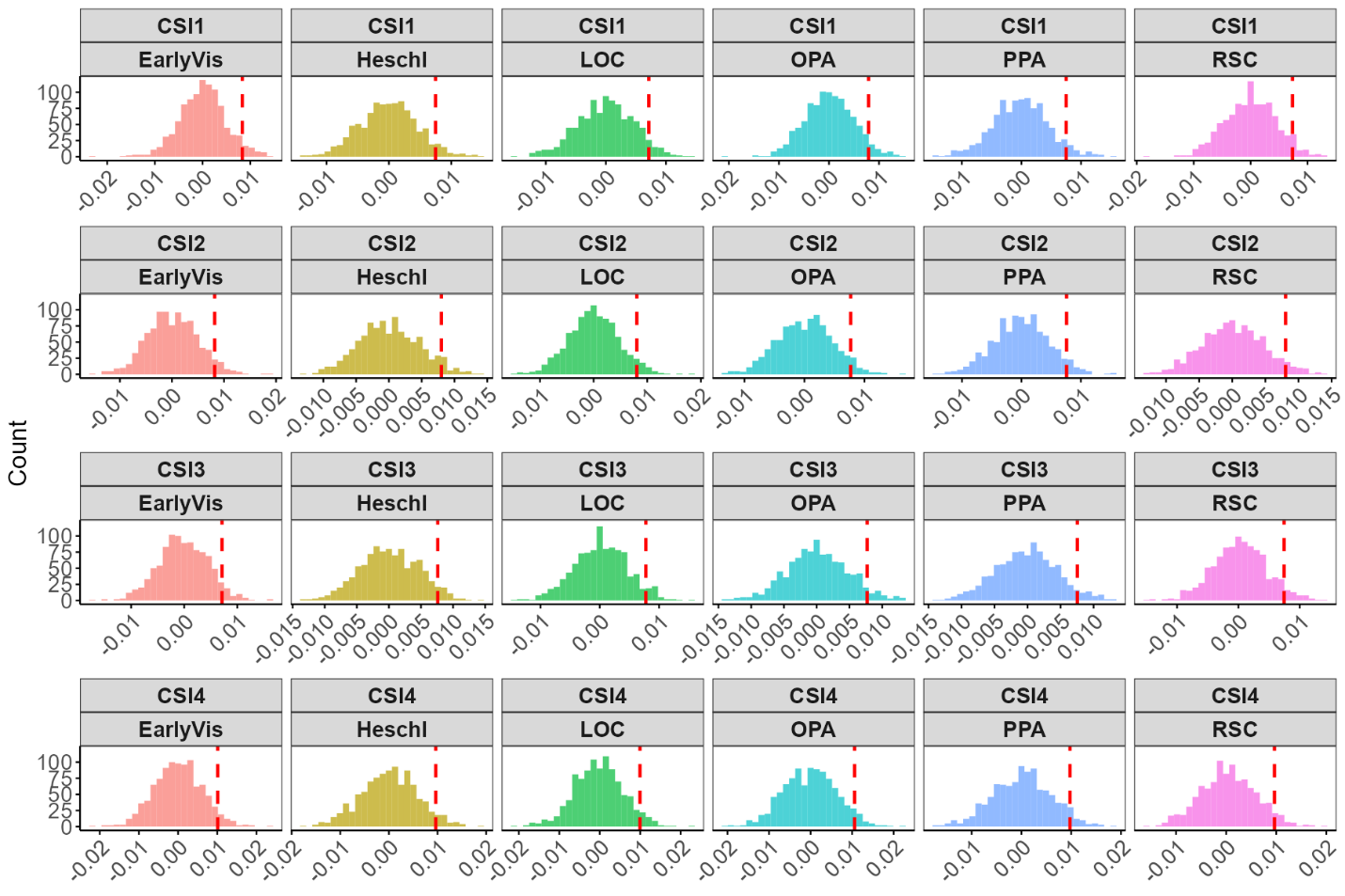

***Fig. S6. Null distributions of correlation coefficients derived from permutations for decoding emotions from fMRI activity in visual cortices.*** *This involved rerunning the Partial Least Squares Regression (PLSR) analyses using the same data and analytic procedure, but with random shuffling of the emotion ratings. This process was iterated 1000 times to generate a null distribution of correlation coefficients (r values). The red line on the x-axis of each subplot indicates the r values where q = 0.05.*

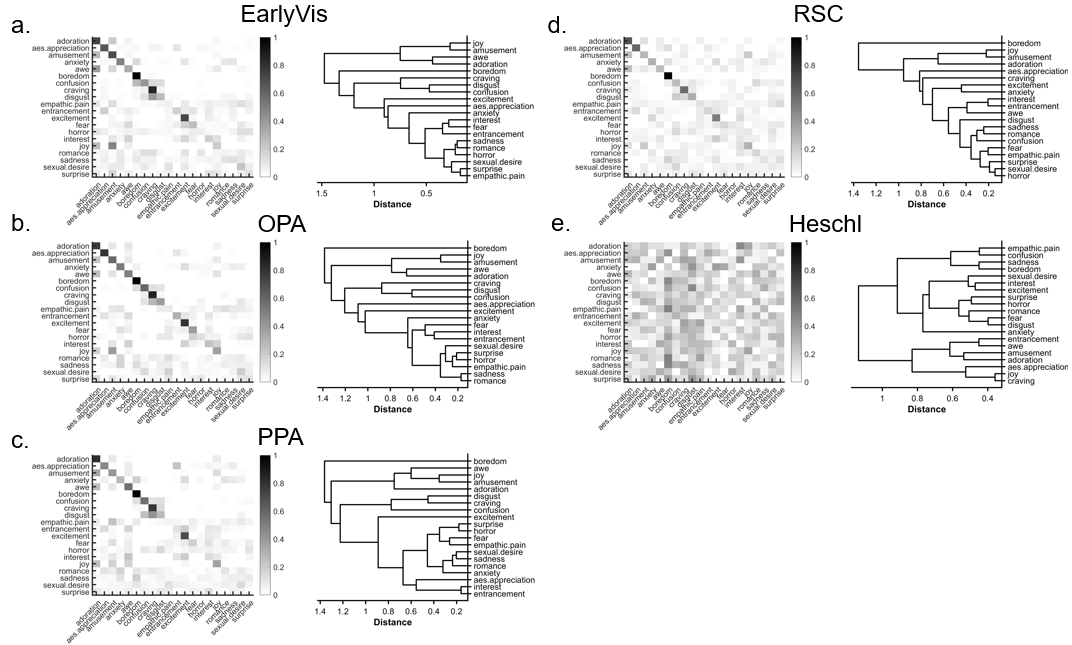

***Fig. S7. The averaged, normalized confusion matrix and dendrogram constructed using Ward's method based on the confusion matrix across four subjects is presented for the relationship between the multivariate pattern responses in different regions including a) Early visual cortex (EarlyVis), b)* *the occipital place area (OPA), c) parahippocampal place area (PPA), d) the retrosplenial complex (RSC), e) Heschl's gyrus (Heschl), and emotion ratings.*** *Rows represent the actual categories of the cross-validated data, and columns denote the predicted categories. Gray colormap indicates the proportion of predictions within the dataset, with each row summing to 1. Correct predictions fall on the diagonal of the matrix, whereas off-diagonal elements reflect wrong predictions.*
